# Modelling individual ampullary afferents in two species of gymnotiform fish using simulation-based inference

**DOI:** 10.64898/2026.06.24.734418

**Authors:** Sarah Mayer, Jan Benda, Jan Grewe

## Abstract

Ampullary electroreceptors are widespread across aquatic vertebrates. The purpose of sensing exogeneous electric fields is conserved across species but the implementations differ and the encoding mechanisms remain incompletely understood. We compared baseline and stimulus-driven response properties of ampullary electroreceptor afferents in the weakly electric fish *Apteronotus leptorhynchus* and *Eigenmannia virescens*. We find that their activity is very well captured by an extended leaky integrate-and-fire model that generalizes across both species. The model shares similarities to a previous model of the tuberous electroreceptor afferents but further incorporates a low-pass pre-filtering and additional noise sources to reproduce the observed spectral response characteristics. The low-pass is essential to shape stimulus encoding in the high-frequency range. Accurate prediction of low-frequency stimulus encoding further requires two distinct noise sources: stimulus-independent white current noise and activity-dependent noise in the adaptation current, which is shaped by the adaptation time constant to yield pink noise dynamics. Using simulation-based inference, we trained a neural network to map model parameters to neuronal response features. This approach enables the generation of heterogeneous, biologically plausible model populations that may serve as a realistic input layer for studying neuronal processing on the next level. With this, we provide a unified and mechanistic model of ampullary electroreceptor encoding in these species and possibly beyond. The proposed model is another step towards a full model of the electrosensory periphery in these animals.

**Author summary:** The ability to sense exogenous electric fields, i.e. passive electroreception, is widespread among aquatic animals. It plays a central role in prey detection and, in some species, also contributes to communication. To understand higher-order brain function, we also need to grasp the sensory periphery and ideally have models that provide naturalistic peripheral responses. Using simulation-based inference (SBI), we here develop a mechanistic model of passive electroreception that is valid for at least two species of electric fish. Through detailed analyses, we identify model components, such as a source of pink noise, that are essential for matching the model’s spectral response properties to those of the recorded cells. The main results of this work are the model itself and the trained inference network (here called the SBI network), which can now be used to create artificial but biologically plausible populations of sensory neurons that may serve as a realistic input layer for studying higher-order neuronal processing. Our work complements the existing models of the active electric sense and is a big step towards a full model of the electrosensory periphery in these animals.

## Introduction

The ability to sense electric fields is widespread among aquatic animals and plays an important role in prey detection. With the so-called passive electric sense, animals can sense exogenous electric fields that are leaked into the water by muscle activity of, for example, prey. Passive electroreception is found in freshwater animals such as catfish, sturgeon, or the mammalian platypus as well as in marine creatures such as sharks and rays [1, 2]. Despite being widespread, the passive electric sense has been lost and re-invented several times during evolution leading to different solutions serving the same purpose (for review e.g. [3, 4]).

Passive electroreception is mediated by primary electroreceptors that are located at the base of cutaneous ampullary organs [4, 5]. Ampullary electroreceptor afferents innervate these organs, encode passive electrosensory information in their spiking activity and carry it to the brain. We here study the response properties of such ampullary afferents in two species of gymnotiform South-American weakly electric fishes, namely the brown-ghost knifefish *Apteronotus leptorhynchus* and the glass knifefish *Eigenmannia virescens* that possess the passive and an active electric sense. These animals actively generate a surrounding electric field through discharges of their electric organ. In contrast to strongly electric fish, which employ their actively generated electric organ discharges for self-defense and prey-capture [6], weakly electric fishes use their actively generated field for prey-detection, navigation [7, 8], and communication [9].

Peripheral encoding as well as central processing of active electrosensory information has been in the focus of research on weakly electric fishes [10] but all fishes that have an active electric sense, are also equipped with a passive electric sense while the converse is not true [3]. Passive electrosensory stimuli, which arise from prey [11, 12], from the animal’s movement within the earth’s magnetic field in sharks [2, 13], or also during courtship signaling in *E. virescens* [14, 15] play an equally important role. The afferents of active and passive sense share some similarities, but there are also profound differences which invite for comparative studies. Ampullary afferents are, as their counterparts of the active system, spontaneously active but show comparatively low levels of intrinsic noise (see [16, 17], for the African mormyrid fish *Gnathonemus petersii* and the South-American gymnotiform fish *A. leptorhynchus*, respectively) and higher levels of nonlinear encoding in the weakly non-linear regime [18].

The afferents of either type project into the hindbrain where those of the active system trifurcate and synapse onto three adjacent maps of the electrosensory lateral line lobe (ELL) while those of the passive system project into a single map, the medial segment [10]. Pyramidal cells in the ELL integrate afferent inputs of populations of afferents [19]. The similarities and also the differences again ask for comparative studies of sensory information processing. A precondition for this, however, is a good grasp of the peripheral encoding properties and, given the current impossibility to simultaneously record from afferent and central neuronal activity, the availability of good models of the input layer. Further, studying processing of passive electrosensory information, we will require good approximations of sensory responses to arbitrary stimuli, not necessarily step- or noise stimuli that may have been used during experimental characterization. In other words, good models for the electrosensory periphery are needed.

Using simulation-based-inference (SBI) [20, 21] we here aim at training an artificial neuronal network that learns the mapping of model parameters to cellular response features. The SBI-network can then be used to retrieve parameter sets that define plausible models of ampullary afferents without the need of a fitting process. The underlying neuron model is based on an extended leaky integrate-and-fire model (LIF, [22–24]) that shares some similarity to the model employed for afferents of the active system [25, 26]. Analysis of the model highlights specific properties of the passive system: we show that two different noise sources are needed to reproduce ampullary response features. One is an additive white current noise in the LIF integrator while the other is part of the activity dependent adaptation process where it is shaped into pink noise through the adaptation time-constant. Further, an additional low-pass pre-filter on the incoming signal is needed to shape the high-frequency decline of the stimulus-response coherence. These specificities are discussed in some detail. Our results further show that the SBI framework could be successfully employed to generate a single SBI-network from which naturalistic models of ampullary cells of at least two different species of gymnotiform electric fish can be constructed. The trained SBI-network allows building arbitrary populations of ampullary cells. Such populations can then be used as input layer for studying subsequent steps of sensory processing which renders it an invaluable tool for future studies.

## Materials and methods

Within this project, we re-evaluated datasets recorded between 2010 and 2012 at the Ludwig Maximilian University (LMU) München. All experimental protocols complied with national and European law and were approved by the Ethics Committees of the Ludwig-Maximilians Universität München (permit no. 55.2-1-54-2531-135-09). The final sample consisted of 34 ampullary electroreceptor afferents (14 in *E. virescens*, 20 in *A. leptorhynchus*) recorded in 9 weakly electric fish of the species *E. virescens* and 14 *A. leptorhynchus*. Electrophysiological recordings were performed on male and female animals. Fish were obtained from a commercial supplier for tropical fish (Aquarium Glaser GmbH, Rodgau, Germany) and kept at a water temperature of 25 °C and a conductivity of around 270 µS cm^−1^ under a 12 h:12 h light-dark cycle.

Before surgery, the animals were deeply anesthetized via bath application of MS 222 (150 mg L^−1^, PharmaQ, Fordingbridge, UK) buffered to pH 7.0 with sodium bicarbonate (150 mg L^−1^, Sigma-Aldrich, Steinheim, Germany). The posterior branch of the anterior lateral line nerve (pALLN) was exposed by making a small cut into the skin covering the nerve. The cut was placed dorsal of the operculum just before the nerve descends towards the anterior lateral line ganglion (ALLNG). Those parts of the skin that were to be cut were locally anesthetized by cutaneous application of liquid lidocaine hydrochloride (2 %, bela-pharm GmbH, Vechta, Germany). During surgery, water supply was ensured through a mouthpiece. To maintain anesthesia, the respiration water contained a dose of 100 mg/l MS 222 again buffered to pH 7.0. After surgery, fish were immobilized by intramuscular injection of 25 µL to 50 µL of tubocurarine (5 mg mL^−1^, Sigma-Aldrich, dissolved in fish saline). Respiration was then switched to normal tank water and the fish was transferred to the experimental tank. In *E. virescens* recordings the electric organ is silenced by the muscle relaxant and was replaced with an artificial electric field that matched in frequency and amplitude to the before-surgery electric field (see [15] for details.)

### Experimental setup

Fish were positioned centrally in the experimental tank, with the major parts of their body submerged into the water. Those body parts that were above the water surface were covered with paper tissue to avoid drying of the skin. Local analgesia was refreshed in intervals of two hours by cutaneous application of Lidocaine around the surgical wounds. Electrodes (borosilicate; 1.5 mm outer diameter; GB150F-8P; Science Products, Hofheim, Germany) were pulled to a resistance of 50 MΩ to 100 MΩ (model P-97; Sutter Instrument, Novato, CA) when filled with 1 mol M KCl solution. Electrodes were fixed in a microdrive (Luigs-Neumann, Ratingen, Germany) and lowered into the nerve. Recordings of electroreceptor afferents were amplified (10x) and lowpass filtered at 10 kHz (SEC-05, npi-electronics, Tamm, Germany, operated in bridge mode). All signals, neuronal recordings, recorded EOD, and the generated stimulus, were digitized with sampling rates of 20 kHz (PCI-6229, National Instruments, Austin, TX). RELACS (www.relacs.net) running on a Linux computer was used for online spike and EOD detection, stimulus generation, and calibration. Recorded data was then stored for offline analysis.

### Identification of ampullary cells

Ampullary cells were identified based on firing rates of 80 Hz to 200 Hz, low levels of the coefficient of variation of the interspike interval distribution of their baseline firing rate, absence of phase-locking to the fish’s EOD, and responses to low-frequency sinusoidal stimuli [17]. We here selected only those cells in which the neuron’s baseline activity as well as the responses to frozen noise stimuli were recorded.

### Electric field recordings

For monitoring the unperturbed EOD, two vertical carbon rods (11 cm long, 8 mm diameter) in a head-tail configuration were placed isopotential to the stimulus. Their signal was differentially amplified with a gain factor between 100 and 500 (depending on the recorded animal) and band-pass filtered (3 to 1500 Hz pass-band, DPA2-FX; npi electronics, Tamm, Germany). As a proxy of the transdermal potential that drives the electroreceptors, we measured the field with two additional silver wires spaced by 1 cm that were located next to the left operculum of the fish and orthogonal to the fish’s longitudinal body axis (amplification 100 to 500 times, band-pass filtered with 3 to 1 500 Hz pass-band, DPA2-FX; npi-electronics, Tamm, Germany). This local EOD measurement recorded the combination of the fish’s own EOD and the applied stimulus.

### Stimulation

Electric stimuli were attenuated (ATN-01M, npi-electronics, Tamm, Germany), isolated from ground (ISO-02V, npi-electronics, Tamm, Germany) and delivered via two horizontal carbon rods (30 cm length, 8 mm diameter) located 15 cm laterally on either side of the fish.

The fish were stimulated with band-limited white noise stimuli with a cut-off frequency of 150 Hz. The stimulus intensity was given as the contrast, i.e. the standard deviation of the white noise stimulus in relation to the fish’s EOD amplitude. The contrast varied between 10 and 20 %. Only cells with at least 20 s of white noise stimulation were included for the analysis.

### Data analysis

Data analysis was done in Python 3 using the packages matplotlib [27], numpy [28], scipy [29], pandas [30], and nixio [31].

#### Baseline analysis

The spontaneous activity of the neurons is given as a sequence of spike times *t*_*i*_ with *n* being the number of spikes. Naturally, the baseline firing rate was calculated as *f*_*base*_ = *n/W* with *W* being the observation interval. From the series of inter-spike intervals (ISI) with the individual intervals given by *T*_*i*_ = *t*_*i*+1_ − *t*_*i*_ we estimated the coefficient of variation of the inter-spike interval distribution (*CV*_*ISI*_ = *σ*(*T*)*/*⟨*T* ⟩) with *T* being the series of intervals, *σ* denoting the standard deviation and ⟨·⟩ denoting averaging. The serial correlation of lag *k* was estimated according to

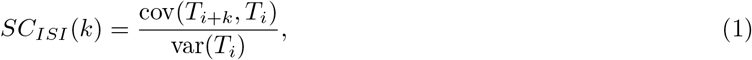

with *cov* and *var* being the covariance and variance, respectively.

#### White noise analysis

In the stimulus-driven case, the neuronal activity of the recorded cell is modulated around the average firing rate that is similar to *f*_*base*_ and in that way encodes the time-course of the stimulus. Spiking activity

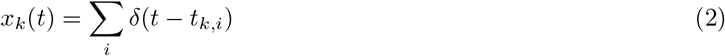

is recorded for each stimulus presentation *k*, as a train of spike times *t*_*k,i*_.

The single-trial firing rate

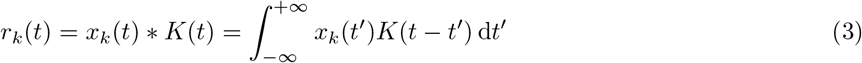

was estimated by convolving the spike train with a Gaussian kernel

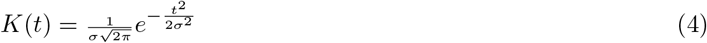

with standard deviation *σ* set to 2.5 ms if not stated otherwise. Averaging over *m* repeated stimulus presentations results in the trial-averaged firing rate

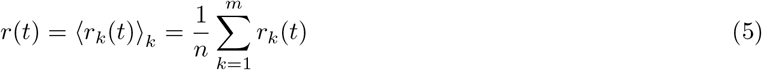

The average firing rate during stimulation, *r*_*s*_ = ⟨*r*(*t*)⟩_*t*_, is given by the temporal average ⟨·⟩_*t*_ over the duration of the stimulus of the trial-averaged firing rate. To quantify how strongly a neuron was driven by the stimulus, we computed the response modulation as the standard deviation 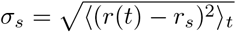 of the trial-averaged firing rate.

#### Spectral analysis

The (model-)neurons are driven by the stimulus and thus their spiking responses depend on the time course of the stimulus. To characterize the relation between stimulus *s*(*t*) and response *x*(*t*), we calculated the first-order susceptibility, also known as the transfer function, and the stimulus-response coherence in the frequency domain.

In the experimental data, stimuli had a duration of 10 s and were cut into segments of 2^14^ samples with 50% overlap resulting in a spectral resolution of 1.22 Hz. Spectra were estimated on individual segments.

The power spectrum of the stimulus *s*(*t*) was estimated as

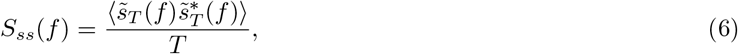

with 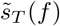 being the Fourier spectrum of the stimulus and 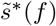 the complex conjugate of 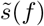 and ⟨·⟩ denoting averaging over segments. The power spectrum of the spike trains *S*_*xx*_(*f*) was estimated accordingly. The cross-spectrum *S*_*xs*_(*f*) between evoked spike trains and stimulus was estimated according to

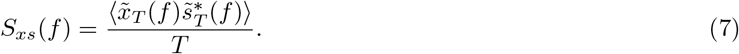

First-order susceptibility, i.e. the transfer function, was then computed from *S*_*xs*_(*f*) and *S*_*ss*_(*f*)

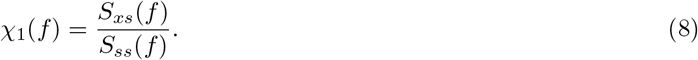

The gain spectrum is the absolute of the complex-valued transfer function

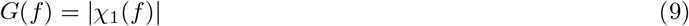

The stimulus-response coherence was estimated according to

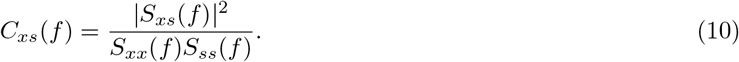

### 0.1 Leaky integrate-and-fire ampullary model

Following the models of the electroreceptor afferents of the active electrosensory system, the P-units [26, 32, 33], we here constructed a leaky integrate-and-fire (LIF) model to reproduce the specific firing properties of ampullary receptors.

First, the input was passed through a low-pass filter that mimics processing in the electroreceptor and dendritic conduction

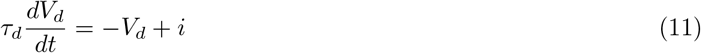

with *V*_*d*_ as the dendritic voltage, *i* the input current, and *τ*_*d*_ being the membrane time constant of the dendrite and the temporal resolution of the simulation (timestep *dt*) was set to 1*s/*20000 = 0.00005*s*, the same as in the experimental recordings.

The dendritic voltage *V*_*d*_(*t*) is then fed into a stochastic leaky integrate-and-fire (LIF) model with adaptation,

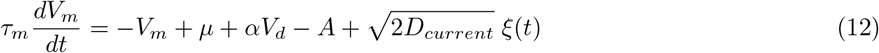

where *τ*_*m*_ is the membrane time-constant, *µ* is a bias current, *α* is a scaling factor for *V*_*d*_, *A* is an inhibiting adaptation current, and 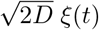 is a white noise with strength *D*_*current*_. Note, that all state variables, membrane voltages *V*_*d*_ and *V*_*m*_ as well as the adaptation current, are dimensionless.

The adaptation current *A* follows

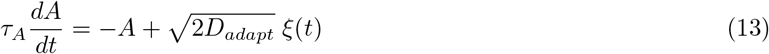

with adaptation time constant *τ*_*A*_ and *D*_*adapt*_ the strength of the adaptation noise.

Whenever the membrane voltage *V*_*m*_(*t*) crosses the spiking threshold *V*_*th*_ = 1, a spike was generated, *V*_*m*_(*t*) was reset to 0, the adaptation current was incremented by Δ*A*, and integration of *V*_*m*_(*t*) was paused for the duration of a refractory period

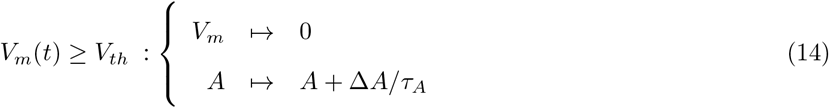

The model is integrated by the Euler forward method with a time-step of Δ*t* = 0.05 ms. The intrinsic noise sources *ξ*(*t*) (Eq 12) for the current and the adaptation noise was added by drawing a random numbers from a normal distribution *N*(0, 1) with zero mean and standard deviation of one in each time step *i*. The noise was then scaled by 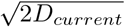 and 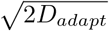, respectively and subsequently divided by 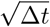. For each trial of a simulation, the variables *A, V*_*d*_ and *V*_*m*_ were drawn from a distribution of initial values and the first second of simulation was discarded as a transient phase.

## Data availability

### Note

The trained SBI network and the cell data will be made freely available on the G-Node data sharing platform gin.g-node.org and receive a DOI upon acceptance of the paper. The trained network is already available for download under https://gin.g-node.org/jgrewe/ampullary_sbi/src/master/packages. The graphical user interface is open-sourced under LGPL and is available on github https://github.com/bendalab/ampullarymodel/.

## Results

The SBI approach consists of an initial brute-force step in which training data is assembled. Sets of model parameters are randomly chosen from prior distributions, the model is run for each parameter set, and respective response features are extracted. Afterwards, an artificial neural network is trained to learn the mapping between model parameters and response features. The first tasks are thus to identify which features are essential to describe the cellular responses and to know their distributions which we used to constrain the priors by excluding implausible simulation results.

### Feature selection from experimental data

Our experimental dataset consists of previously collected data from ampullary afferents recorded in *E. virescens* and *A. leptorhynchus* [15, 17]. The selection of cells used here is constrained by the availability of sufficiently long recordings of spontaneous activity and responses to noise stimuli at a given intensity (20% relative to fish’s self-generated field, see methods). This leaves us with a sample of 14 and 20 cells, respectively. For these, we extracted response features as depicted for single-cell examples in Fig 2 and S1 Fig.

**Fig 1.**
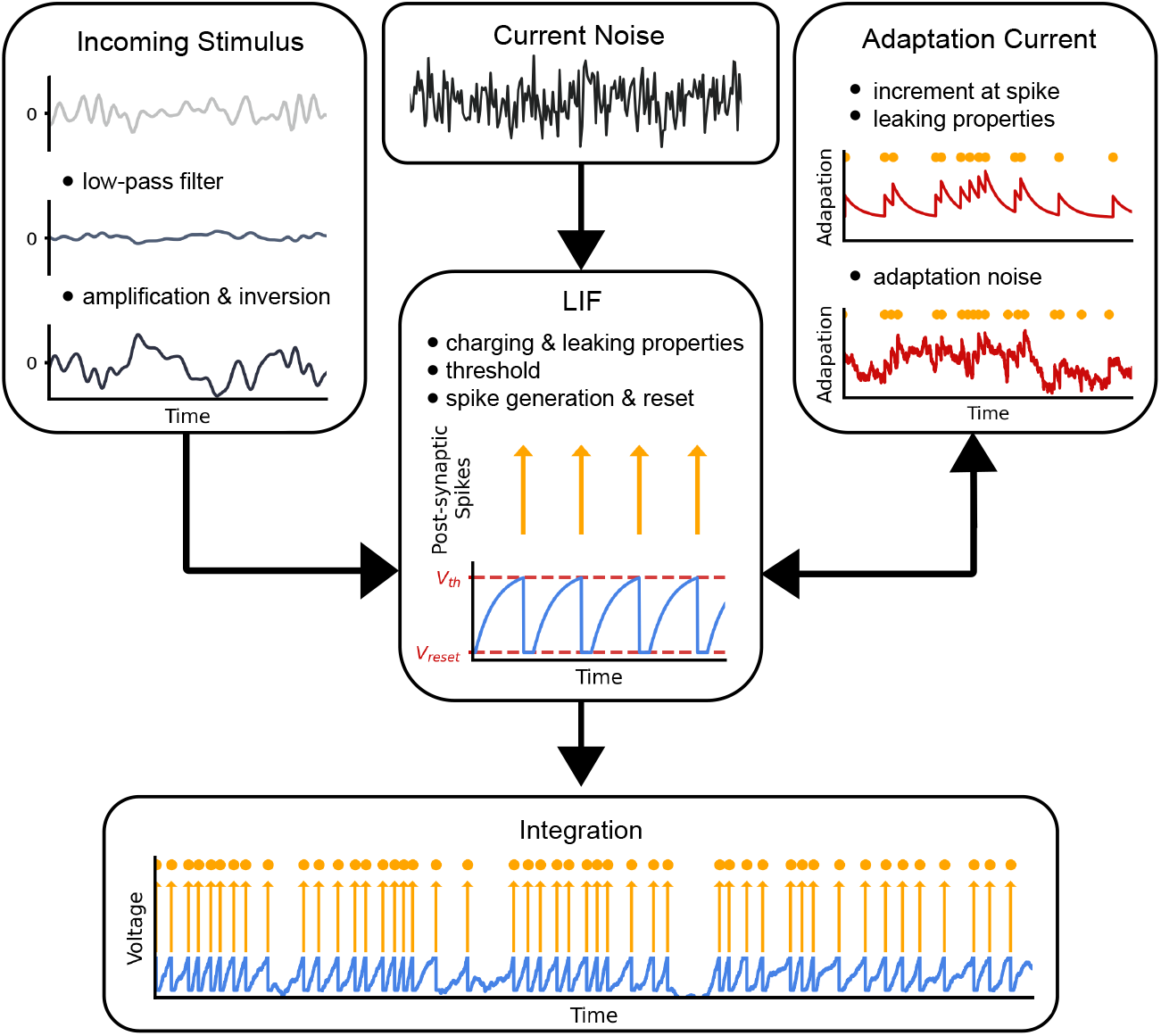
Sketch of the ampullary model. Ampullary afferents are modeled by an extended leaky integrate-and-fire model (LIF, see methods for details). The core is the LIF integrator that integrates the membrane voltage and generates action potentials. The model input is the external stimulus (*s*(*t*)) that is low-pass filtered (*τ*_*dendrite*_) and scaled (*α*) before it is sent into the integrator (left box). An additional white current noise current with the noise strength (*ξ*_*current*_, top box) is added. With each action potential (orange dots), an inhibitory adaptation current is incremented (*δ*_*adapt*_) that relaxes back to zero with a given time-constant (*τ*_*adapt*_) and is subjected to additional white noise (*ξ*_*adapt*_, right box). The LIF core (center box) integrates these inputs. Once a given threshold (V_*th*_) is exceeded, the time of the action potential (orange arrows and dots) is noted, the adaptation incremented, and the voltage is reset to the reset voltage (V_*reset*_). Integration is paused for a certain refractory period.

**Fig 2.**
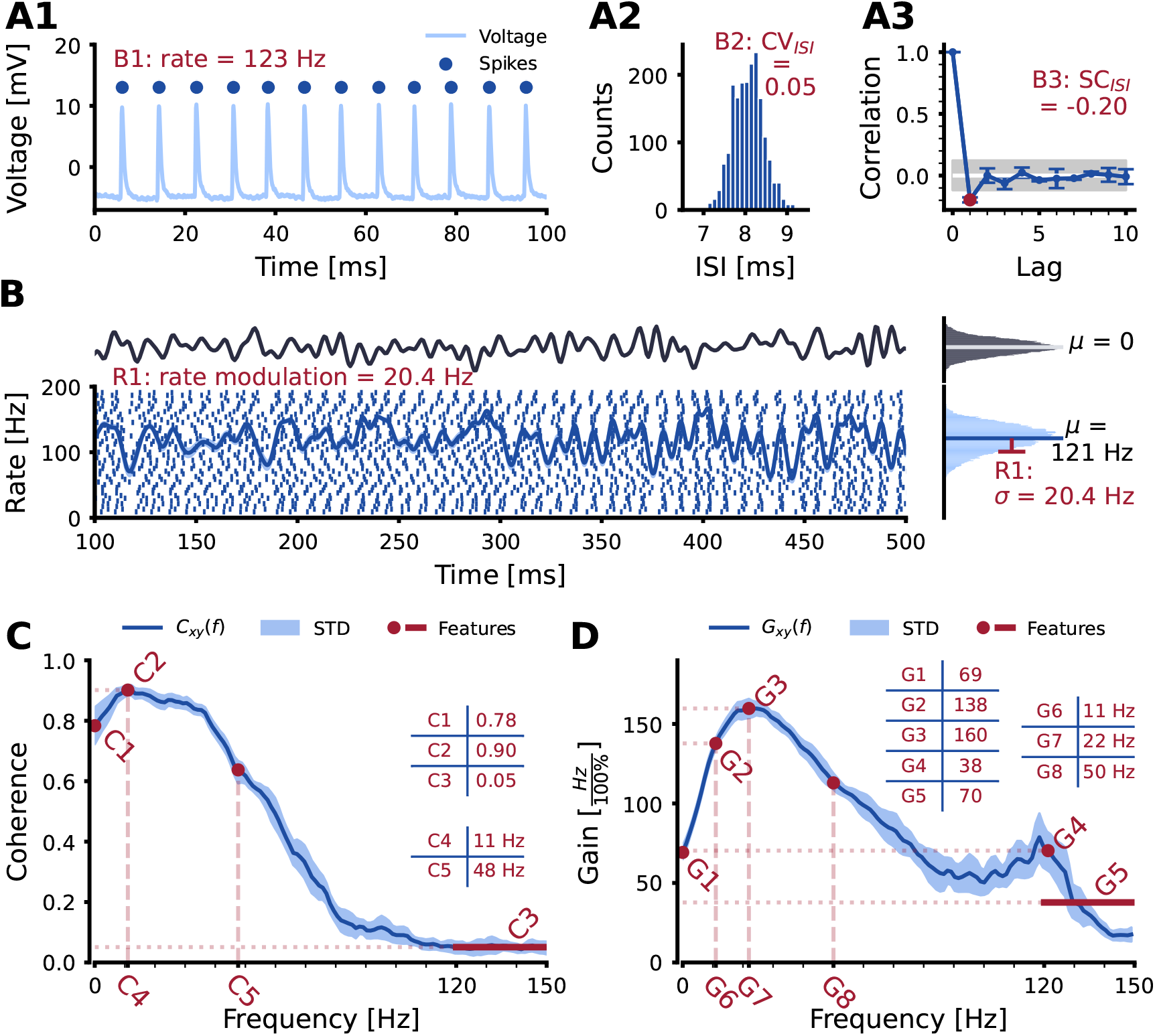
Baseline and stimulus-driven responses of an exemplary ampullary afferent recorded in *Apteronotus leptorhynchus*. **A1-3** Characteristics of the spontaneous baseline response from which the baseline features (*B1 – B3*, see text) were extracted. **A1** Short section of the recorded voltage trace, dots indicate detected action potentials. **A2** Distribution of the interspike intervals of the spontaneous activity. **A3** Serial correlation of the interspike intervals. **B** Stimulus driven response of the same cell. The neuron was stimulated with a frozen zero–mean band–limited white noise stimulus with spectral content up to 150 Hz (top trace, see methods for details). Bottom plot shows the spiking response to repeated presentations of the same stimulus together with the average firing rate estimated by kernel-convolution with a 2.5 ms Gaussian kernel. The shaded area around the firing rate shows the SEM across trials. Histogram on the right shows the firing rate distribution, which describes the strength of the response (feature *R1*). **C** Stimulus-response coherence. Solid line shows the across-trial average and the shaded area the across-trial variability. The shape of the coherence spectrum was captured by the five highlighted features *C1 – C5*. **D** The gain of the transfer function. Again, the solid is the across-trial average and the shaded area the across-trial variability. Eight features (*G1 – G8*) were used to characterize the shape of the transfer function.

#### Baseline activity

From the spontaneous baseline activity, we extracted three characterizing features: *B1: baseline rate*, the average firing rate in the absence of a driving stimulus, *B2: CV*_*ISI*_, the coefficient of variation of the interspike interval distribution that describes the response regularity of the spontaneous activity, *B3: SC*_*ISI*_, lag-1 serial correlation of the interspike intervals that characterizes the dependence of successive spikes (Fig 2 panels A1 – A3).

#### Stimulus–driven activity

Our experimental dataset contains recordings under stimulation with frozen band-limited noise stimuli. The stimulus had a zero mean and spectral power in the range 0 Hz to 150 Hz. The stimulus intensity was calibrated to amount to 20 % of the fish’s self-generated field. Fig 2 B shows the respective neuronal responses in form of spike-rasters and the firing rate estimated via kernel-convolution with a Gaussian kernel (*σ* = 2.5 ms). From the noise responses we extracted a single feature describing the sensitivity of the neuron (*R1:* rate modulation) that is calculated as the standard deviation of the firing rates observed under stimulation (histogram on the right).

The responses to dynamic stimuli were further analyzed by means of the stimulus-response coherence and the transfer function [34]. The coherence describes the linear correlation between stimulus and responses and is a spectral measure varying between zero, no linear correlation, and one, perfect linear correlation. Deviations from unity are caused by noise, non-linear behavior of the system, or combinations thereof. The shape of the coherence spectrum was characterized by five features: *C1: C*_0*Hz*_, the coherence at zero frequency; *C2: C*_*max*_, the maximum coherence; 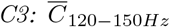, the average coherence at high frequencies (120 – 150 Hz); *C4: F*_*max*_, the frequency of the maximum coherence; and *C5: F*_*cutoff*_, the cutoff frequency of the coherence spectrum.

The transfer function (or first-order susceptibility) is the ratio of the cross-spectrum of stimulus and response and the stimulus’ power-spectrum. Its absolute value, the gain spectrum, describes the spectral sensitivity of the system. The shape of the gain spectrum was captured by eight features: *G1* : *G*_0*Hz*_, the gain at zero frequency; *G2:* 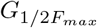, the gain half-way between zero and maximum; *G3: G*_*max*_, the maximum gain; *G4: G*_*rate*_, the gain at average firing rate which, in weakly driven cells, can show a distinct peak; *G5: G*_120−150*Hz*_, the average gain at frequencies in the range 120 – 150 Hz; *G6:* 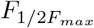 the position of the gain halfway to the frequency of the maximum of the gain; *G7: F*_*max*_, the position of the maximum gain; and *G8: F*_*cutoff*_, the cutoff-frequency of the gain spectrum. Feature *G6* depends only on the position of the maximum and is redundant. We still kept it to emphasize the initial part of the transfer function during training.

### Similar feature distributions with differences

We made use of the so-called *RestrictionEstimation* feature of the SBI package that restricts the prior to regions more likely to yield valid simulations (e.g. avoiding parameter settings leading to unrealistically high or low firing rates). We could thereby speed up the simulation phase by increasing the likeliness of successful models. To constrain this network we first need to know the feature ranges of the experimental dataset that are shown in Fig 3.

**Fig 3.**
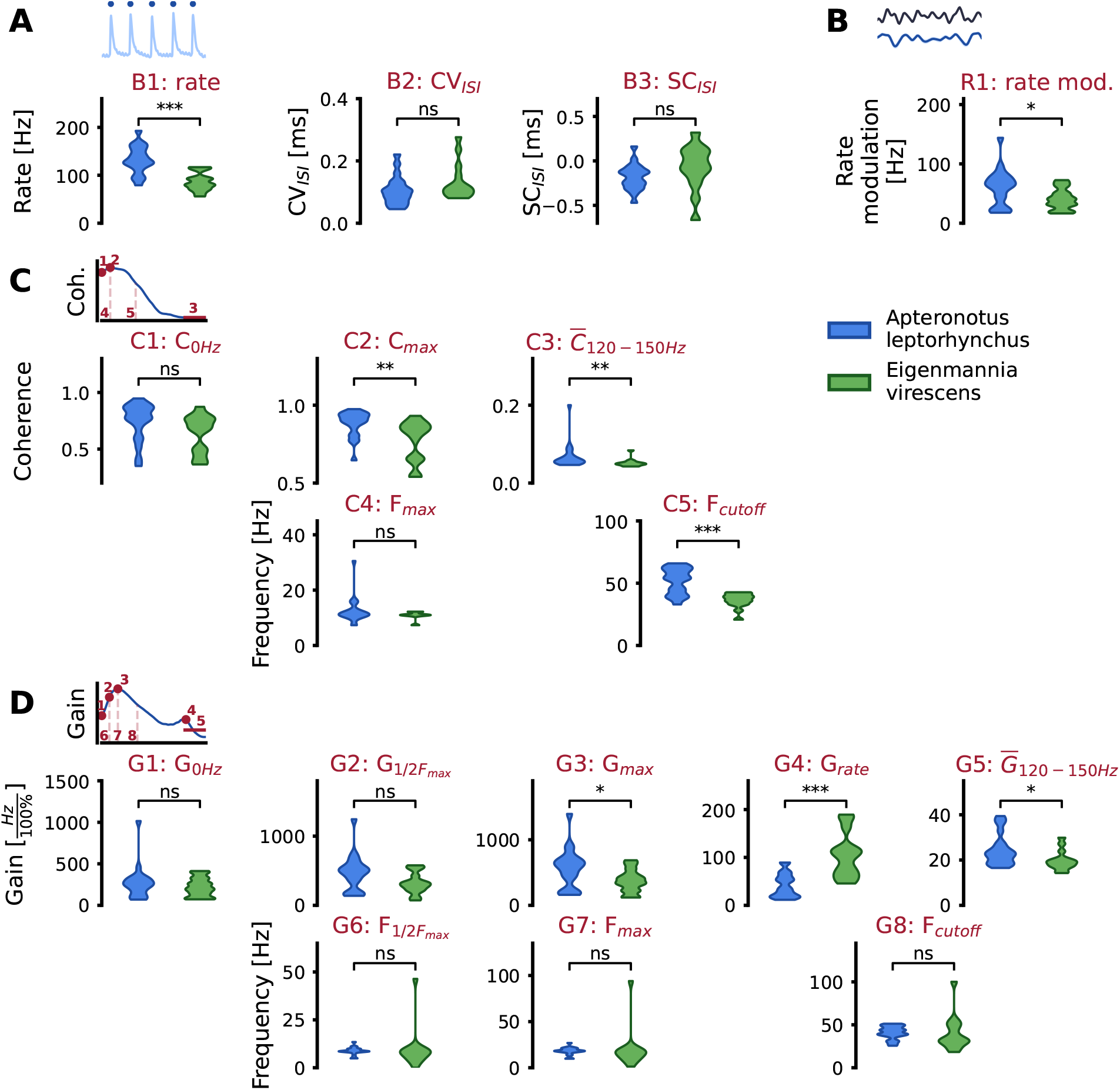
Distributions of the response features of cells recorded in *A. leptorhynchus* (blue) and *E. virescens* (green) **A** Baseline features *B1 – B3*. **B** Distribution of the rate modulations in the stimulus-driven response (*R1*). **C** Characteristics of the coherence spectrum. Features *C1 – C3* are coherence values while features *C4, C5* are the positions on the frequency axis. Horizontally aligned subplots in the top and bottom row describe the same point on the spectrum. **D** Characteristics of the gain spectrum. Top row features *G1 – G5* are given in the unit of the transfer function, while those in the bottom row (*G6 – G8*) describe the positions on the frequency axis.

At a first glance, it appears that ampullary afferents in *E. virescens* and *A. leptorhynchus* are very similar and indeed most feature distributions are not statistically significantly different. There are, however, some differences: The average baseline firing rate (*B1*) is higher in *A. leptorhynchus* while the *CV*_*ISI*_ (*B2*) is slightly lower (Fig 3A). An elevated average firing rate may then explain why ampullary cells in *A. leptorhynchus* have slightly increased rate modulation capabilities when driven by a stimulus (Fig 3 B, feature (*R1*). An increased firing rate further affects the cutoff frequency of the coherence function (*C5*) [35]) and the slightly higher peak coherence (*C2*) may result from less noise, smaller *CV*_*ISI*_. Features describing the gain are very similarly distributed, only the maximum gain (*G3*) and the gain at baseline firing rate (*G4*) are different. A higher maximum gain goes along with higher rate modulations in *A. leptorhynchus* (*R1*) and this also affects feature *G4* as this peak is only prominent in cells that are only weakly driven by the stimulus [17].

### One fits all: Successful reproduction of cellular data for both species

We then trained the SBI-network on a feature-set based on 12 million simulations using the final model with model parameters that were randomly chosen from prior distributions (see discussion of some detailed model considerations). Once the SBI-network is trained, it can be queried to return the parameter set that will most likely match the experimentally observed features best. For this, we applied the MAP (maximum-a-posteriori) approach provided by the SBI package: i.e. multiple samples are drawn from the trained posterior and used as starting points for a gradient-ascent optimization that searches for highest posterior density. Fig 4 compares experimental responses and simulation results obtained from the MAP estimate for the same *A. leptorhynchus* cell as in Fig 2. The same SBI-network was also queried to provide parameter sets to reproduce *E. virescens* ampullary cells. Even though the distribution of the same features were different in the samples we used here, the SBI-network still returns parameter sets that reproduce neuronal responses of either species similarly well (S2 Fig). Note that this involves no actual fitting to experimental data. It is only the vector of desired response features of baseline and stimulus-driven responses that is used to draw parameter sets from the SBI-network. Baseline features match almost perfectly. Also, responses to white-noise stimuli are very nicely reproduced. There are, however, also some parts of the responses that are not so well captured (between 400 and 450 ms in Fig 4 B or at ms 260 in S2 Fig B for an *E. virescens* ampullary afferent). The stimulus-response coherence as well as the gain spectrum are very well-matched in both example cells.

**Fig 4.**
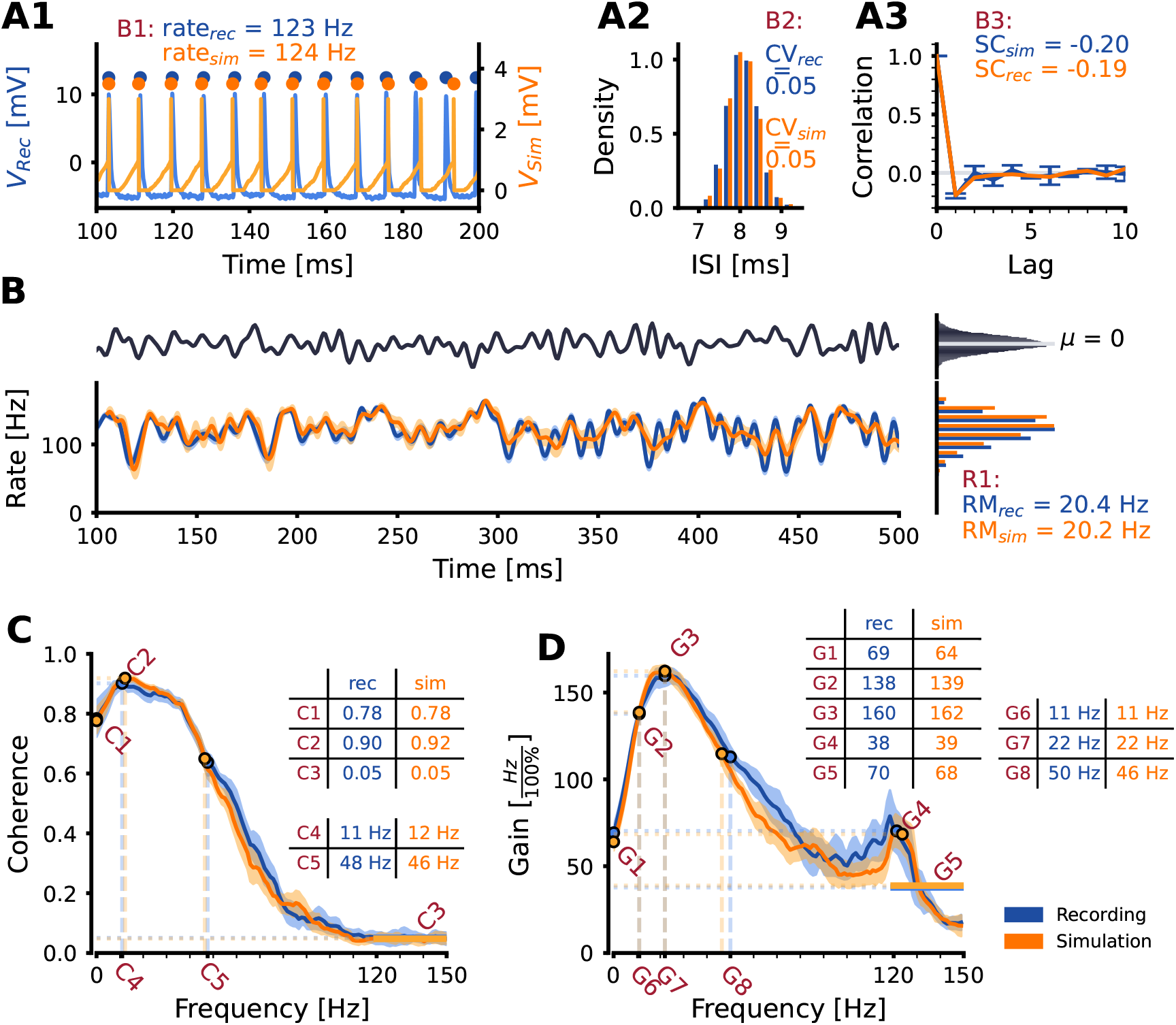
Comparison of experimental data and model predictions. Response features of an exemplary ampullary afferent recorded in *A. leptorhynchus* (blue) and the simulated counterpart (orange). **A1-3** The characteristics of the spontaneous activity are well reproduced. Note that the waveform of the simulated action potentials in A1 is a sketch and not part of the actual model output. **B** Depicts the stimulus-driven response to the frozen noise stimulus. The rate modulations, i.e. the standard deviation of the histograms on the right match nicely. Shaded areas show the SEM across trials. Note that the delay between cellular response and simulated response is arbitrary and was adjusted for best match between recording and simulation. **C, D** Stimulus-response coherence and gain spectra. Table insets compare the extracted features of the recordings and the simulations. Shaded areas around the trial-averaged spectra are the standard deviation across trials.

Over all cells, the trends we have seen in the example data are conserved. Features of baseline and stimulus-driven responses are almost perfectly matched (Fig 5 A & B). The characteristics of the stimulus-response coherence are well-matched, but the simulations tend to overestimate the coherence at zero frequency (*C1*), the peak of the coherence (*C2*), and the coherence at high frequencies (*C3*). The position of the peak (*C4*) and the position of the cutoff frequency (*C5*) are slightly underestimated. Correlations between recorded and simulated data are sometimes reduced because of the limited spectral resolution which is particularly relevant for the position of the maximum (Fig 5 C). The features of the gain function, however, are generally well-matched. Only the gain at high frequencies (*G5*) seems to be slightly overestimated by the simulations. Goodness of fit and deviations are similar for ampullary afferents of either species. S3 Fig offers a slightly different view on the comparison of measured and simulated response features. Most cellular response features are captured nicely by the simulation (horizontal lines). It seems that some of the more extreme feature values cannot be reproduced by the models drawn from the SBI-network. We observe both cases: extreme values in the recordings are “forced” towards the mean of the distribution (e.g. the lowest peak coherence values in S3 Fig C or extremely low feature values for *G6 & G7* in D) or they become even more extreme in the simulation (e.g. *C3* in S3 Fig). Extreme features seem harder to capture than “well-behaving” cells. One must note, however that features *C3* and *G5*, i.e. the coherence respectively gain at high frequencies are somewhat sensitive as these measures are strongly affected if the firing rate is in the range 120-150 Hz. Some features are generally more easy to reproduce than others. S4 Fig shows distributions of difference between z-scores for each feature in recordings and simulations. Some features are easier to faithfully reproduce than others. The serial correlation (*B3*), for example, shows rather wide ranges of z-score differences while the baseline response of the unperturbed spontaneous activity or the response modulation in the stimulus-driven response are very well reproduced, features of the coherence show higher prediction errors. The gain of the transfer function, on the other hand, appears mostly easy to reproduce. For some features this can be directly understood: the *G1: G*_0*Hz*_ is linked to the average firing rate, and the maximum gain (*G3: G*_*max*_) is related to the cell’s sensitivity and quantifies a similar response property as the rate modulation (feature *R1*). There is, however, no obvious difference between species and model responses match real data very well.

**Fig 5.**
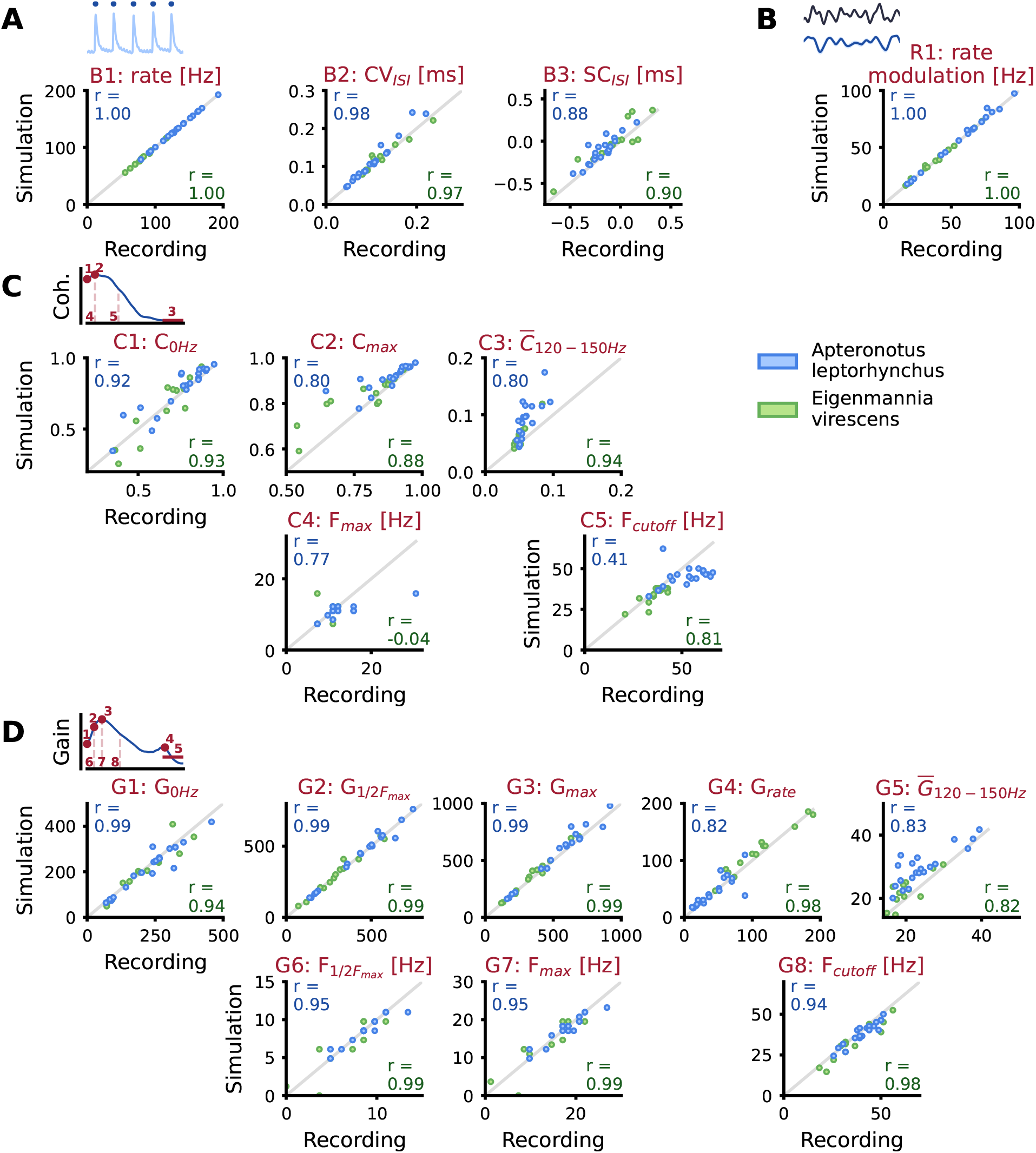
Comparison of response features in recordings and simulations for *A. leptorhynchus* and *E. virescens*. Data for *A. leptorhynchus* are shown in blue and those of *E. virescens* are plotted in green. The respective Pearson correlation coefficients are given in the top-left and bottom-right corner, respectively. Ideally, data should be on the diagonal. **A** The three baseline features. **B** The stimulus-driven firing rate modulation. **C** Comparison of features describing the stimulus-response coherence. **D** Comparison of the features describing the gain spectrum.

### Model parameters are not constrained by the prior distributions

The training dataset is constructed from parameter sets randomly drawn from prior distributions of the respective parameters. These prior distributions were manually constrained by plausibility considerations and prior knowledge gathered in test runs. The refractory period, for example, was limited to the range 0 – 3 ms which seems plausible considering maximum firing rates that were observed experimentally. The distributions in Fig 6 represent the parameters chosen by the SBI-network to reproduce experimentally recorded data. Y-axes depict the respective ranges of the prior distributions. With few exceptions, parameter distributions are not constrained by prior limits. Medians are well within the ranges and do not accumulate at the borders. Only a few parameters (offset current *I*_*offset*_ and input scaling *g*_*stimulu*_) are statistically significantly different between the two species which must be taken with care since the number of cells we could use for our approach is limited. A higher offset current *I*_*offset*_ in *A. leptorhynchus* seems to go along with higher baseline firing rates seen in Fig 3 A. Further, some parameters may be linked and depend on each other. This can now be studied using the trained SBI-network to draw models with similar features and analyze the chosen parameter sets. This, however is beyond the scope of the present work.

**Fig 6.**
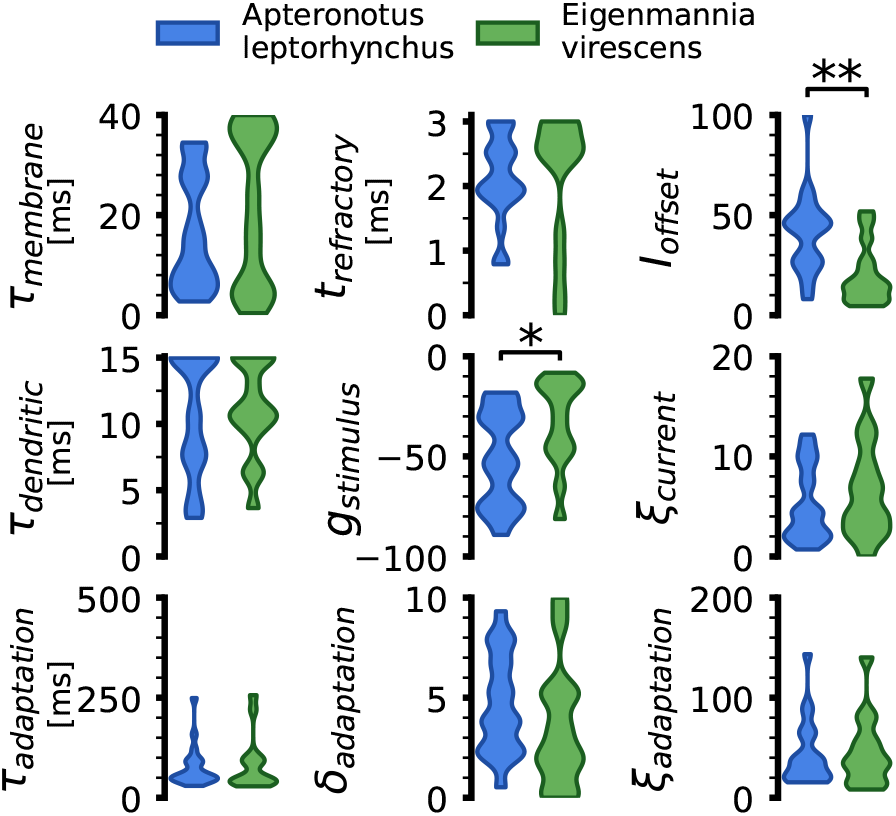
Comparison of model parameters for *A. leptorhynchus* (blue, n=20) and *E. virescens* (green, n=14). The range on the y-axis reflects the range of the prior distributions. Parameter distributions were tested on statistically significant differences using a Mann-Whitney U-Test. Statistical significance was denoted with asterisks (∗: p *<* 0.05, ∗∗: p *<* 0.01).

### Model generalizes well for dynamic stimuli

We challenged the model’s generalization power by reproducing the responses to novel stimuli such as a band limited white noise stimulus of lesser intensity and step stimuli that were not used to train the SBI network. Responses to these stimuli were only available for a subset of the cells in our training set.

Figure Fig 7 compares the responses to weaker stimuli for two example cells for each species (panel A, left: *A. leptorhynchus* and *E. virescens* on the right). Generally, simulated (orange) and recorded (blue and green, respectively) responses match well for most parts. There are, however, sections in which the model prediction deviates (Fig 7 A, right, between 850 and 900 ms). For across-cell comparison we here pooled the data from both species and compare the stimulus-driven features *R1:* rate modulation (Fig 7 B), root-mean-square (RMS) distances between the recorded and simulated coherence spectra (Fig 7 C), and RMS distance between the recorded and stimulated gain spectra (Fig 7 D) at both stimulus intensities. The rate modulation is also very well captured for the low-intensity stimulus, the Pearson correlation coefficient between recording and simulation at 10% contrast is still very high (0.97 vs. 0.99 for 20%contrast). The prediction quality for the coherence spectra is equally good for 20% and 10% contrasts, the average error amounts to only 5% of the coherence. The prediction of the gain spectrum is slightly worse for the untrained contrast of 10% (statistically significant with p *<* 0.001, Wilcoxon signed-rank test for paired samples). S5 Fig shows recorded and simulated examples of gain spectra for both species in which the peak at baseline firing rate is very pronounced at 10% contrast while it is almost not visible at the higher contrast that was used to sample the model parameters from the SBI network. The relatively large RMS distances arise from a slight underestimation of the gain for frequencies below the firing rate.

**Fig 7.**
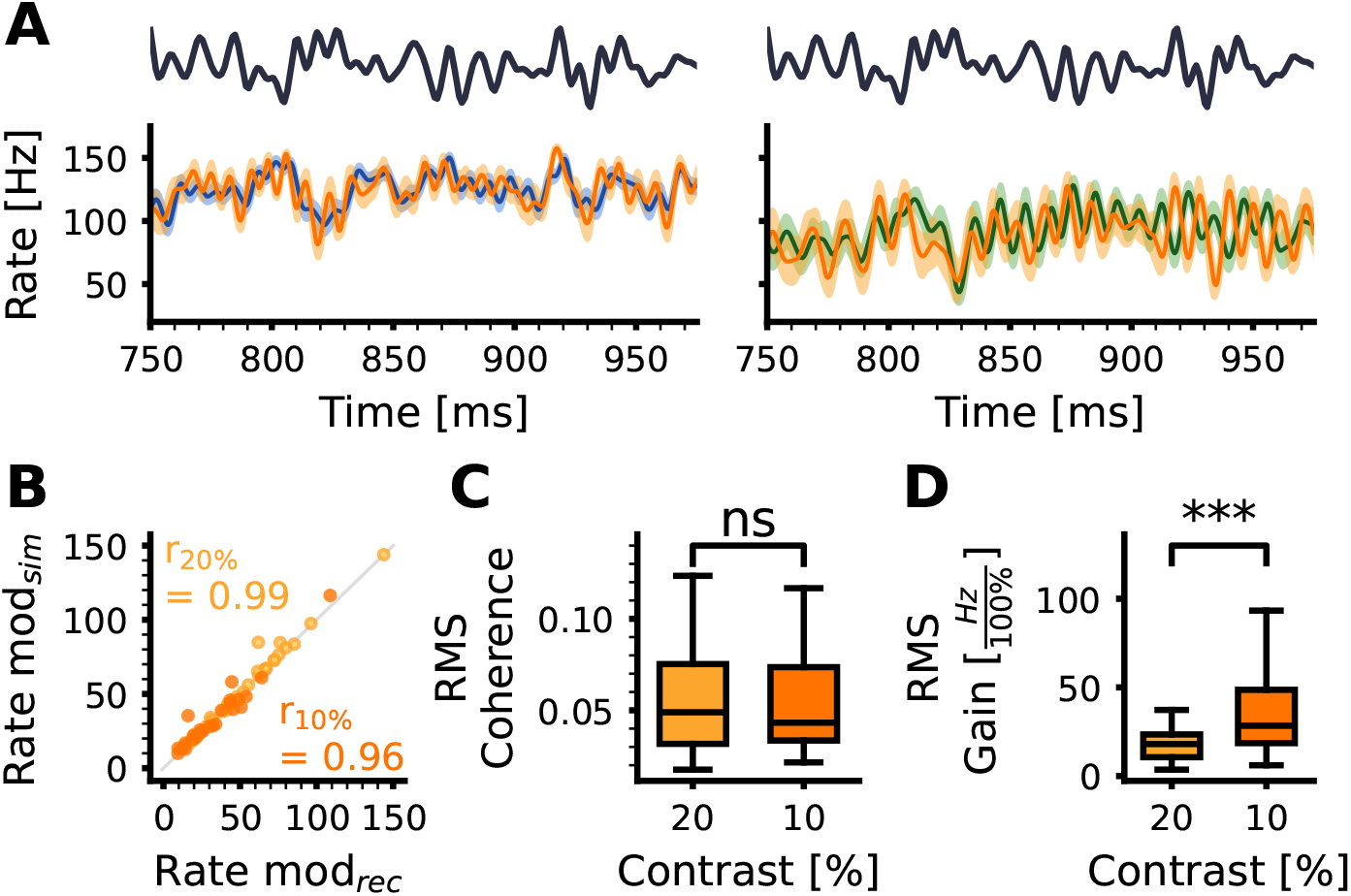
Test for model generalization using non-trained white-noise stimulus. In the training process, we extracted features from neuronal responses to a noise stimulus with a specific intensity (20% contrast). In subsets of our experimental data we also recorded responses to weaker noise stimuli. We used these recordings to challenge our model. That is, we asked the trained SBI-network for model parameters reproducing the 20% contrast features and then simulated with a stimulus of a different intensity. **A** two example responses to 10% contrast stimuli for *A. leptorhynchus* and *E. virescens* (left and right panel, respectively.) **B** comparison of the rate modulations. Light orange are the data points for 20% and dark orange for 10% contrast stimuli. We here pooled data from both species (14 *A. leptorhynchus* and 12 *E. virescens*). **C** Root-mean-square distance between simulated and recorded coherence spectra at 20% (trained) and 10% contrast (untrained) stimuli. Statistical significance was tested using a Wilcoxon signed rank test. **D** Same as C but for the gain spectrum.

For a different subset of cells we additionally recorded responses to step stimuli. Even though step-responses were not part of the training procedure, the simulated cells reproduce the onset responses to step stimuli nicely for either species S6 Fig.

## Discussion

In the present account we have applied simulation-based inference (SBI) [21] to develop a model for ampullary electroreceptor afferents in gymnotiform electric fish. In the SBI approach, an artificial neural network is trained to learn the mapping between model parameters and response features, there is no actual fitting involved. The resulting SBI-network can be queried to provide model parameterization that will likely lead to the desired features. We show that SBI can be successfully applied to reproduce responses of ampullary afferents in two species of South American weakly electric fish. With its availability, we lay important groundwork for studies on the integration of passive electrosensory information on higher processing steps. It becomes possible to create large populations of biologically plausible individual model neurons without going through a fitting process that provides specific solutions for individual previously recorded neurons (see also S2 Appendix).

To successfully apply SBI, some preconditions must be met: (i) the model must be powerful enough to actually reproduce experimentally observed cellular responses and (ii) the selected response features must be effective descriptors of the neuronal response properties. In what follows, we will first discuss these two issues before we turn to the generalization capabilities of the model.

### Model considerations

The chosen model is based on a leaky integrate-and-fire model (LIF, Fig 1) inspired by models used to describe p-type afferents of the tuberous electroreceptor pathway (P-units) [18, 26, 32, 36]. Tuberous and ampullary signal pathways consist of similar building blocks: (i) primary electroreceptors are located at the base of cutaneous pits that differ in shape and signal transduction properties; (ii) primary electroreceptors are connected to respective afferent fibers via synapses in the form of large boutons or calical structures (ampullary and tuberous organs, respectively) [37]; and (iii) afferent fibers send action potentials along their axons to the ELL in the hindbrain. Accordingly, the models share some similarities but there are also differences. Ampullary cells encode the direct electrosensory input, not amplitude modulations of the self-generated field that P-units encode; hence, there is no non-linear amplitude-modulation extraction mechanism involved. In the following paragraphs we will highlight similarities and differences between the models.

#### Activity-dependent adaptation current

Much like P-units, ampullary afferents show pronounced spike-frequency adaptation in response to step stimuli (S6 Fig), i.e. cells respond initially strongly to a step and then rapidly adapt towards a steady-state response level. This is here modeled as an inhibitory adaptation current: i.e. with each fired action potential, the adaptation current is incremented and decays with a certain time-constant. This is the same implementation as in models by [26, 38]. Chacron et al. [32] yield a similar effect by transiently incrementing the spiking threshold in their model. Adaptation currents and dynamic thresholds, however, have distinctly different effects. While inhibitory adaptation currents act linearly and shift the neuron’s firing-frequency vs. stimulus intensity (*f* − *I*) curve, the dynamic threshold acts divisively and changes the slope of the *f* − *I* curve [39]. The shift induced by inhibitory adaptation current matches the observed effects of different adaptation levels on the *f* − *I* curves of P-units [40]. Inhibitory adaptation effectively acts as a high-pass filter which leading to a drop in the gain for low-frequencies [38]. Our dataset does not allow for an equivalent analysis but there is also no indication suggesting a different adaptation process. Rather, gain and coherence spectra can be reproduced nicely even when stimuli of different intensities, contrasts, are used. We thus assume that adaptation in ampullary cells follows the same rules as in the tuberous electrosensory pathway.

#### Additional (dendritic) low-pass filter to attenuate high-frequency components

In the P-unit model, an incoming signal is passed through a low-pass filter that is considered a representative of passive conduction in pre- and postsynaptic regions at the synapse between primary electroreceptors and afferents. In that model, low-pass filtering is needed to adjust spike-time locking to the self-generated field. Further, the signal is passed through a cubic threshold non-linearity required to extract amplitude modulations of the self-generated field [18, 26]. While the latter is not required in ampullary afferents, the incoming signal is also low-pass filtered before it serves as input to the LIF spike generator.

The low-pass filter again mimics temporal properties of the primary electroreceptor cells (which were shown to be low-pass in the catfish *Ictalurus nebulosus* [41]), passive signal conduction, and synaptic properties. Indeed, it is essential to reproduce high-frequency cutoffs of the stimulus-response coherence in both species (Fig 8). Gain and coherence spectra of recorded data (blue and green data for *A. leptorhynchus* and *E. virescens*, respectively) show high-frequency cutoffs well below 100 Hz and the coherence spectrum, in particular, shows a rather steep fall-off for frequencies exceeding 50 Hz. These spectra of both species are well-matched by the full model (orange). Removing the (dendritic) low-pass filter, however, has strong effects on gain and coherence spectra: The stimulus-response coherence now shows elevated coherence for frequencies in the range 60 – 120 Hz that is not seen in recorded data. The membrane time constant (*τ*_*m*_) in the integrator is the second low-pass filter which acts on the sub-threshold membrane voltage. Increasing *τ*_*m*_ in an attempt to compensate for the lacking dendritic filter does not suffice to shape the coherence spectrum (not shown).

**Fig 8.**
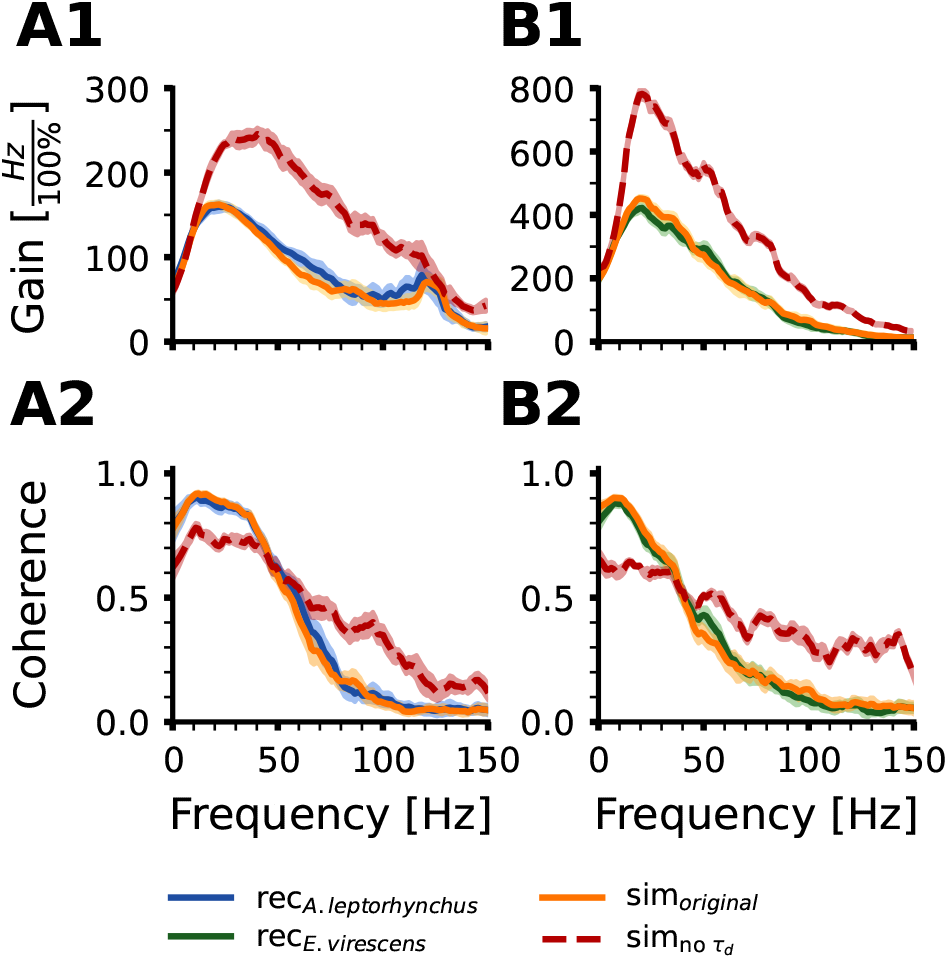
High-frequency cutoff requires an additional low-pass filter. **A1** Gain spectrum of the transfer function of an exemplary ampullary afferent recorded in *A. leptorhynchus* (blue) and the best matching model (orange). The red broken line shows the gain spectrum of a model that does not contain the additional low-pass filter but is parameterized in the same way as the original model. **A2** The coherence spectrum for the same configurations as in A1. **B1, B2** The same for an exemplary cell from *E. virescens*.

#### Coherence function is shaped by two noise sources

Transfer function and coherence spectrum of ampullary afferents in both species show a distinct drop-off for low frequencies (*<* 20 Hz, Fig 2 C & D). As discussed above, attenuated low-frequency susceptibility, is due to the adaptation current. The drop in low-frequency coherence, on the other hand, can not be explained by inhibitory adaptation currents [42] but is either due to increased non-linearity or an impaired signal-to-noise ratio in low frequencies.

Fig 9 shows the impact of different noise sources on the stimulus-response coherence. Without any noise, condition *ξ*_0_, the coherence is highest and does not show a low-frequency drop. With only a white-noise current *ξ*_*c*_ in the LIF spike generator, the peak coherence can be successfully adjusted. Still, *ξ*_*c*_ is not sufficient to reproduce the desired drop at low frequencies (yellow and orange lines in Fig 9). Adding noise only to the adaptation process (*ξ*_*adapt*_, red line in Fig 9) leads to a low-frequency drop, but the overall coherence is too high. *ξ*_*adapt*_ is low-pass filtered by the adaptation time-constant *τ*_*adapt*_ which converts it to pink noise [42]. Combining pink noise of the adaptation and white noise in the LIF core, finally yields good matches between experimental data and model responses which is similar to the P-unit model suggested by Chacron and colleagues [32]. The noise in the adaptation process is turned into pink noise through the adaptation time-constant which attenuates high-frequency power. This is slightly different to the Chacron model [32] who explicitly used two Ornstein-Uhlenbeck processes with different time-constants to model correlations that are induced for example by quantal transmitter release at synapses [43]. Our simulations, however, show that we need LIF-intrinsic white noise and noise in the adaptation current to obtain the best match between experimental and simulation data (see also S1 Appendix). The need for two noise sources is in line with findings in the locust auditory receptors, where it was concluded that one source of colored noise originates from adaptation currents and interactions with the adaptation time-constant [42, 44].

**Fig 9.**
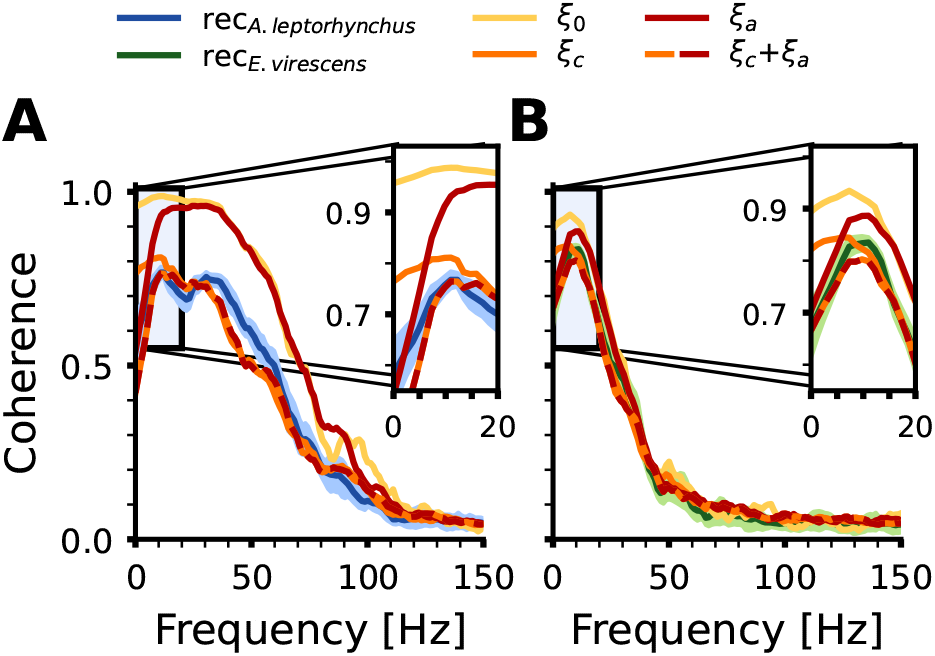
Two noise sources are required to shape the coherence function correctly. **A** Stimulus response coherence of an example *A. leptorhynchus* cell (blue solid line) and of the best fitting models that contain no noise (yellow) only additive white noise in the spike generator (orange), only additive noise in the adaptation (red) or both noise sources (hatched red-orange line). The inset highlights the critical low-frequency range. **B** The same for an exemplary cell from *E. virescens*.

### Feature selection

To train the SBI-network we defined a set of features that were extracted from spontaneous baseline responses as well as responses to band-limited white noise stimuli. The data was gathered in two species of South-American weakly electric fishes, namely the brown ghost knifefish *Apteronotus leptorhynchus* and the glass knifefish *Eigenmannia virescens*, which we recorded from in different contexts [15, 17]. Of all cells reported in the respective original papers, only a subset could be re-used in this study as only these were stimulated using the same stimuli.

The choice of features may influence the final parametrization of the respective model cells. To create our set of models for the active, tuberous, electroreceptor afferents, P-units [18, 26, 45], the respective model was fitted to cells in which no white-noise stimulus was available but rather responses to step-stimuli were used to estimate *f* − *I* curves. These models reproduce white-noise responses well and also show the theoretically expected non-linear behaviors in the weakly-nonlinear regime [18, 46]. Some parameters, such as the dendritic low-pass filter that is also part of our ampullary model, is not well constrained by step-stimuli [35]. On the other hand, in this study, we did not consider step stimuli but focussed on those datasets that contained white-noise responses as dynamic stimulation is closer to natural stimulus situations [12] and noise stimulation allows for system’s characterization with much less recording time. On the downside, it may underestimate non-linearities due to linearizing effects of noise [18, 47, 48]. The models we retrieved from the SBI-network, though based on white noise data, can reproduce step responses quite well even though there were no features describing them (S6 Fig).

A linear system can be fully characterized by measuring its responses to white noise stimuli from which the transfer function (and the absolute of it, the gain) as well as the coherence function can be estimated. Multiplication of transfer function and the stimulus Fourier spectrum allows to predict the system’s overall responsiveness which we here also characterize with the response modulation. Deviations of the coherence function from unity is in parts attributable to internal noise which we also estimate by the CV of the interspike interval distribution under baseline conditions. Thus, there are features that provide overlapping information, or may correlate. For the SBI process this is not a problem, since the SBI-network will learn the interdependencies and would even learn to ignore uninformative features [20, 21]. Thus, we would argue to rather have more than too little features. A drawback of this approach is that the retrieval of model parameters requires to provide desired values for all features. In the accompanying graphical tool (S2 Appendix), the SBI-network can be used to retrieve model parameters and the tool assists by providing feature distribution from the training set that can be constrained by the user.

### SBI approach does not require extensive datasets

Experimental data provides the ground truth of modelling approaches. Yet, experimental data can be flawed as well and rigorous data curation is necessary. The curation processes applied here were hurtful as quite substantial amounts of recorded cells needed to be excluded for various reasons: (i) too little data to faithfully estimate the white noise responses, as they have not been the focus of the *E. virescens* recordings made in context of encoding of electrocommunication signals [15], (ii) instability during the recording of the baseline activity which usually is done in the first 30+ seconds after a recording could be established, or (iii) exclusion due to wrong stimulus conditions. In the end, our work is based on 14 cells recorded in *E. virescens* and 20 cells recorded in *A. leptorhynchus* which may seem not too impressive. For the SBI approach this is not critical: Experimental data is essential to establish the feature space and to learn whether features effectively constrain the model parameters. For the training set used to train the SBI-network, however, experimental data are of no relevance as it does not fit but learns how model parameters map to response features. Accordingly, the experimental data serve as tests to see if the parameters retrieved from the trained network actually lead to plausible results. As we have demonstrated, this is satisfactory; the important factor here is that the experimental example data are good representatives of the overall variety of cells we studied.

### Prediction power and applicability across species

Even though the ability to sense exogenous electric fields is widespread and apparently conserved across species, this does not mean that ampullary electroreceptors are the same across species. On the contrary, passive electroreception has been lost and regained several times. Although these systems serve the same purpose, they differ with respect to response modes and morphology [4, 49]. In teleosts, these receptors are characterized by responses to anodal stimuli, i.e. firing rates increase when the outside becomes more positive relative to the inside. In contrast, their counterparts in non-teleost fish, including paddlefish, skates, and lampreys, increase firing in response to stimuli of the opposite sign [4]. What these systems share is a mechanosensory origin, and primary electroreceptors are connected to afferents via ribbon synapses [49]. They are all clearly low-frequency tuned and show spontaneous activity in the absence of any driving input. Spontaneous firing rates in the catfish *Ictalurus nebulosus* were found in the range 35 - 60 Hz [50] with a clear temperature dependence [51], in the range 45±7.5 Hz in the skate *Raja eglanteria* [52], 55±8.5 Hz reported for the paddlefish *Polydon spatula* [12]. In the African weakly electric fish *Gnathonemus petersii* we see higher firing rates in the range 73±23.8 Hz [16] which fits well to the range we here describe for *E. virescens* ampullary cells but is lower than we find for *A. leptorhynchus* (Fig 3 [17]). Spectral sensitivity scales along with firing rates in the sense that cells with higher firing rates encode stimuli in a wider range than those with lower firing rates, which is a generic feature of spiking neurons [35]. The question arises whether the model suggested here is of general applicability for ampullary afferents in other species. From inspections of the data published on ampullary afferents of *G. petersii* [16] we are very optimistic as all response features fall into the feature ranges we find in our experimental data and are covered by the training set. For the paddlefish, however, we are pessimistic, as the response spectra of these show the two oscillations [12] that we touched upon above and that cannot be reproduced by our model. Our model deviates from the model of paddlefish ampullary electroreceptors [12]. There, two oscillators are identified, the first oscillates at the spontaneous firing rate of the afferent and a second epitheliar oscillation of the ampullary canal. Our model only contains oscillation at the spontaneous firing rate which is implicitly created by interaction of offset current and membrane time-constant. As a consequence, our model will have limited generalization power for ampullary encoding in paddlefish and others which show two oscillations. The spectral sensitivity of primary electroreceptors in catfish was modelled with sequences of low-pass filters [41] which fits well to the dendritic low-pass filter applied here and, to our knowledge, no oscillations are described in the literature, which is promising. For the skate, we are lacking the required data. It would be nice to perform comparative studies to see how far the model carries across species.

## Conclusions

With the present study we developed a phenomenological LIF-based model of ampullary electroreceptor afferents. The model is powerful enough to reproduce all response features that we observe in electrophysiological recordings made in two species of weakly electric fish, namely *Eigenmannia virescens* and *Apteronotus leptorhynchus*. We further provide descriptions and population statistics for these cells. Using the SBI framework we trained an artificial neuronal network, that has learned the mapping between model parameter and response features and can now be used to create ampullary model cells with desired response features. This is an invaluable tool for example to create populations of neurons that respects the level of heterogeneity of natural neuronal populations and thus fosters future studies.

## Acknowledgements

Within this project we re-evaluated data that were recorded by Henriette Walz, Anna Stöckl, and JG. We are further grateful to Benjamin Lindner for discussions in early stages of the project and Jakob Macke for discussions and help regarding the SBI package.

## Supporting information

**S1 Fig.**
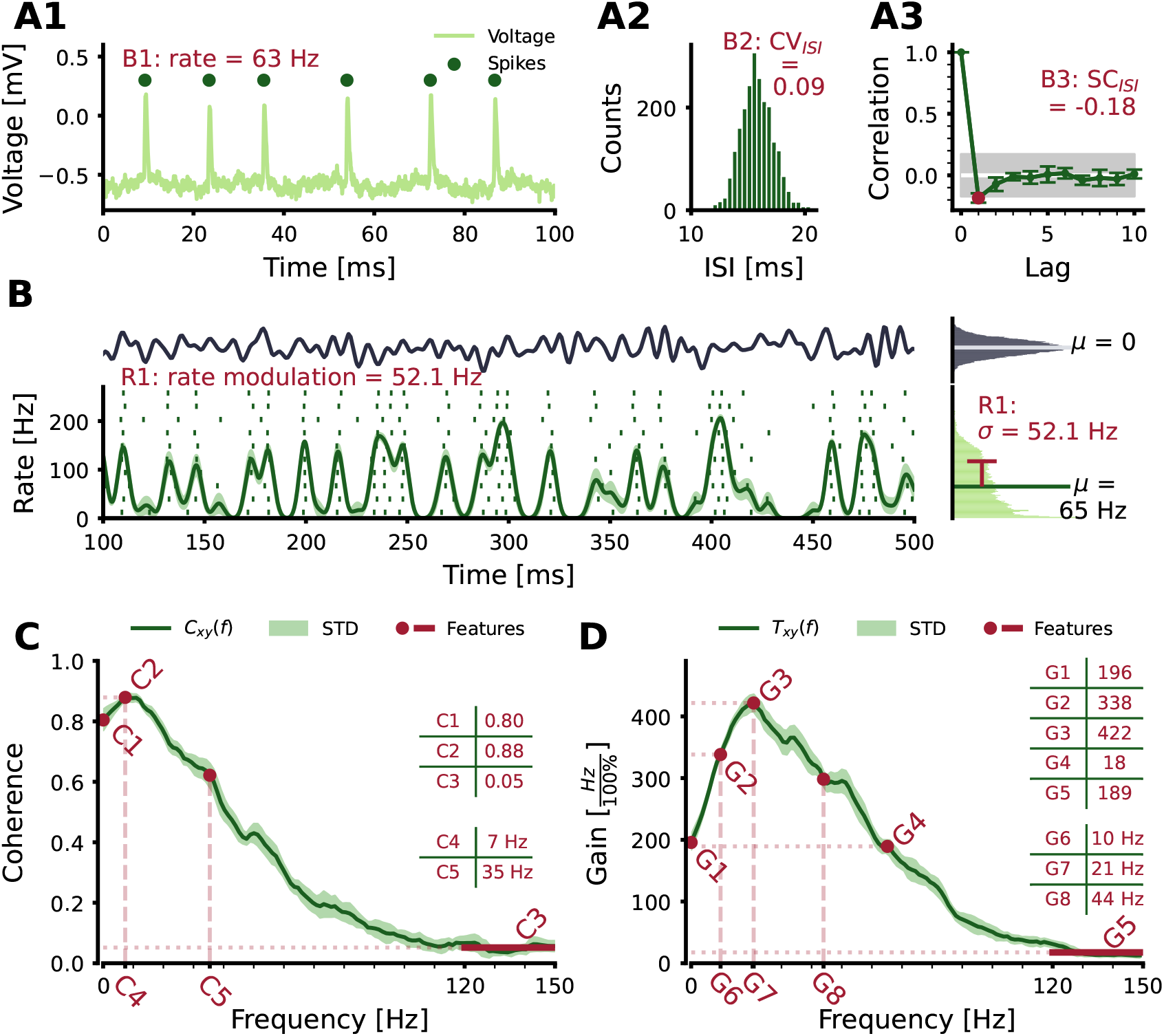
Response features of an Eigenmannia ampullary afferent. Same as Fig 2.

**S2 Fig.**
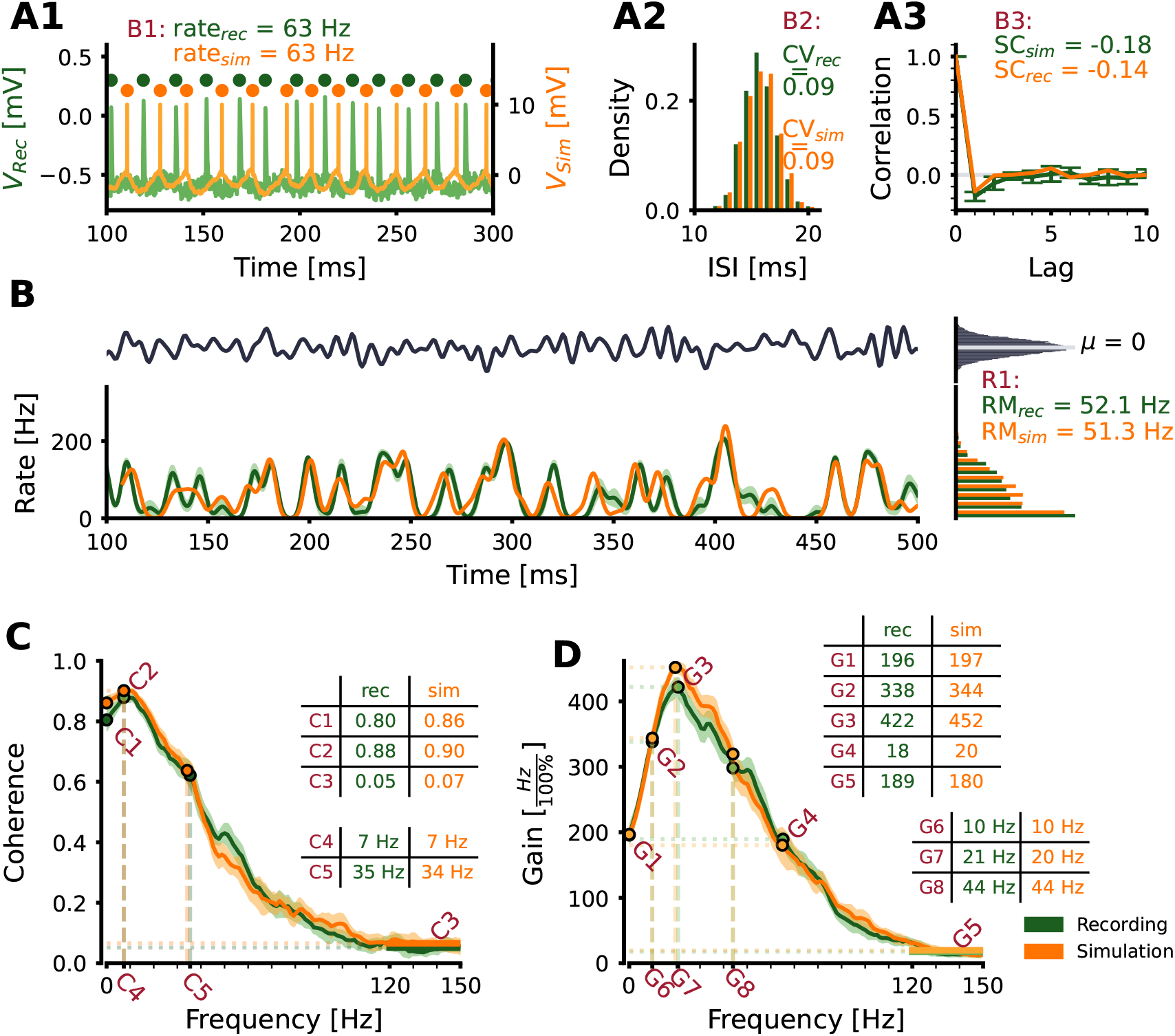
Comparison of *E. virescens* ampullary responses and model predictions. Same as Fig 4 but for an ampullary cell recorded in *E. virescens*.

**S3 Fig.**
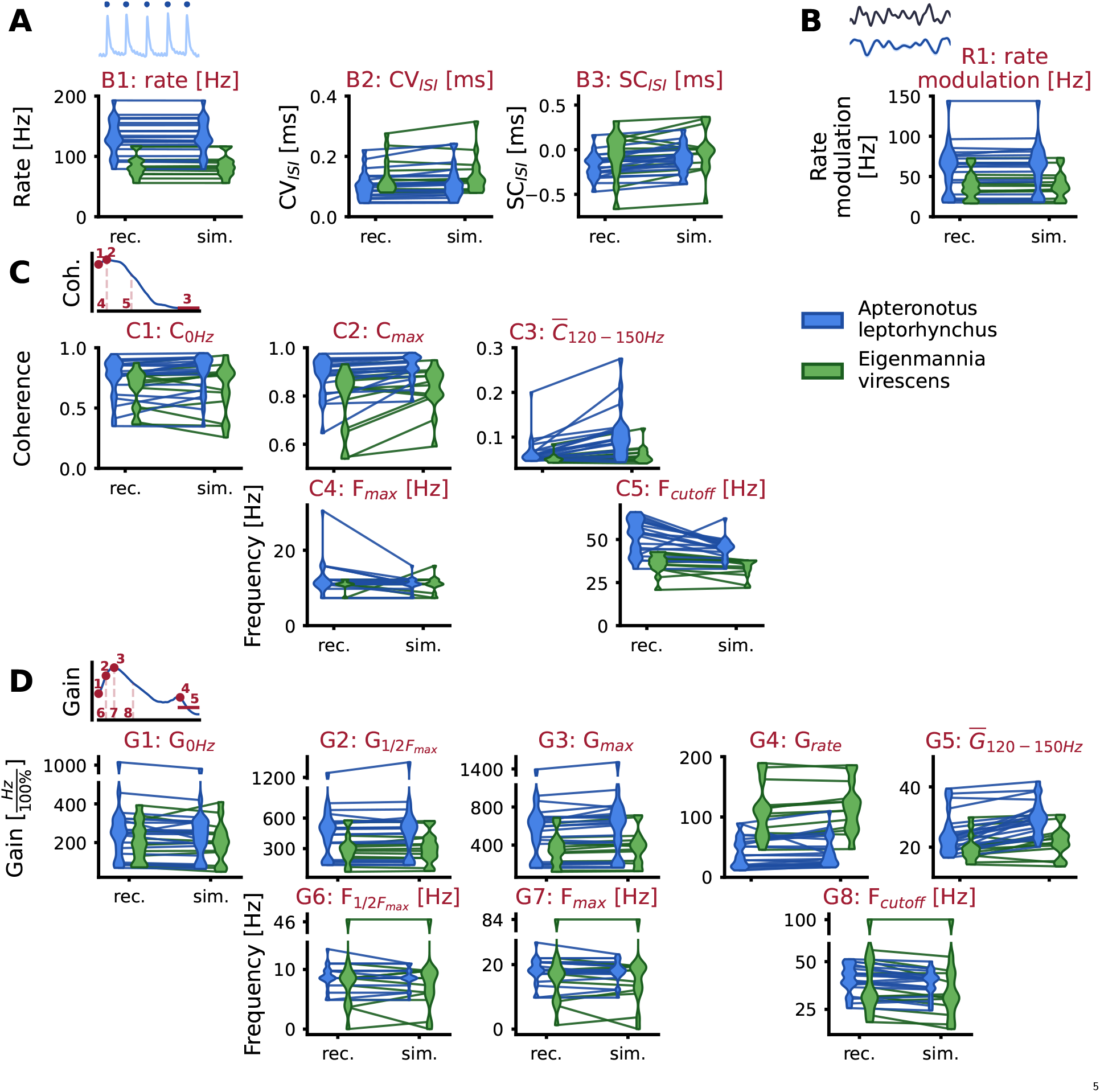
Cell-by-cell comparison of recorded and simulated features. Violin plots show distributions of experimentally recorded (left) and simulated response features (right) for both species. Lines connect corresponding real and simulated data.

**S4 Fig.**
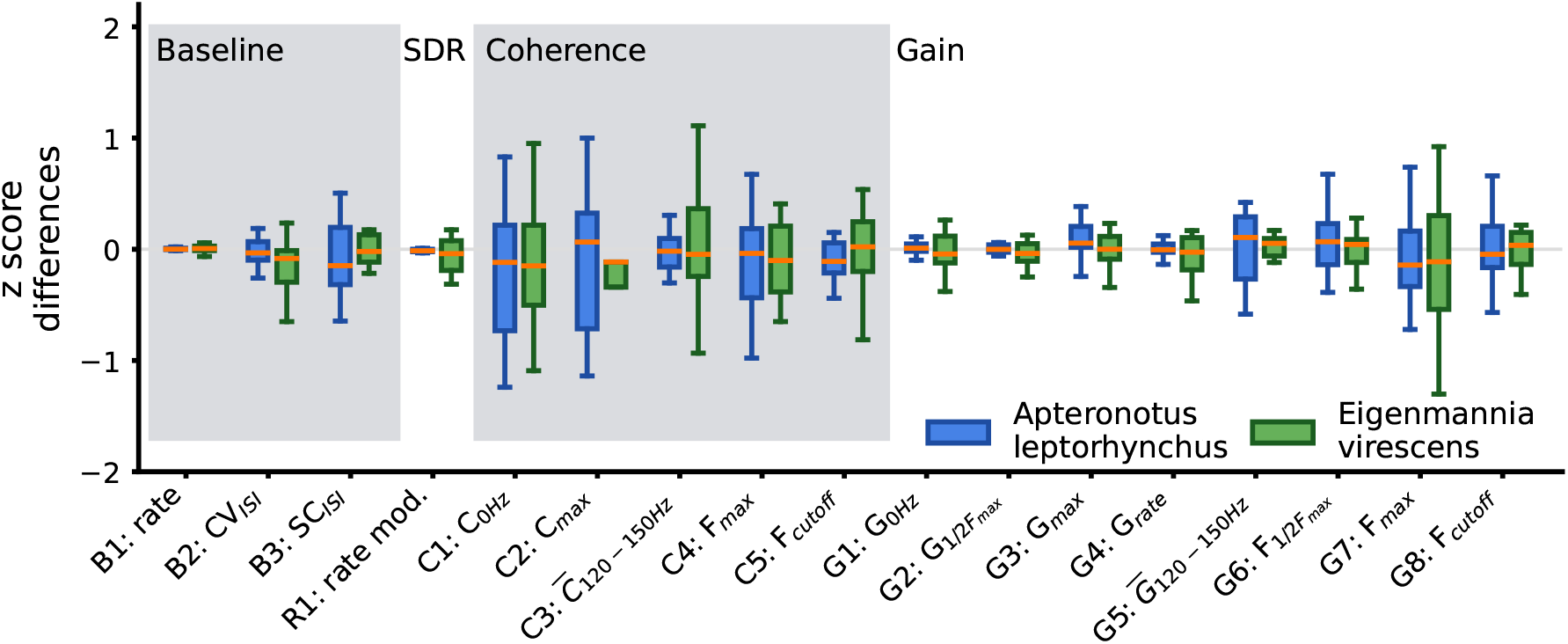
Easy vs. difficult features. The simulations can reproduce different features with different ease. For each feature we compare the differences in z-scores between recording and simulation for both species. Wide distributions indicate that a feature is harder to reproduce than a feature which shows a narrow distribution. A median that is below or above the zero line, would indicate a systematic under- or overestimation of the given feature. The abbreviation SDR stands for “stimulus-driven-response”.

**S5 Fig.**
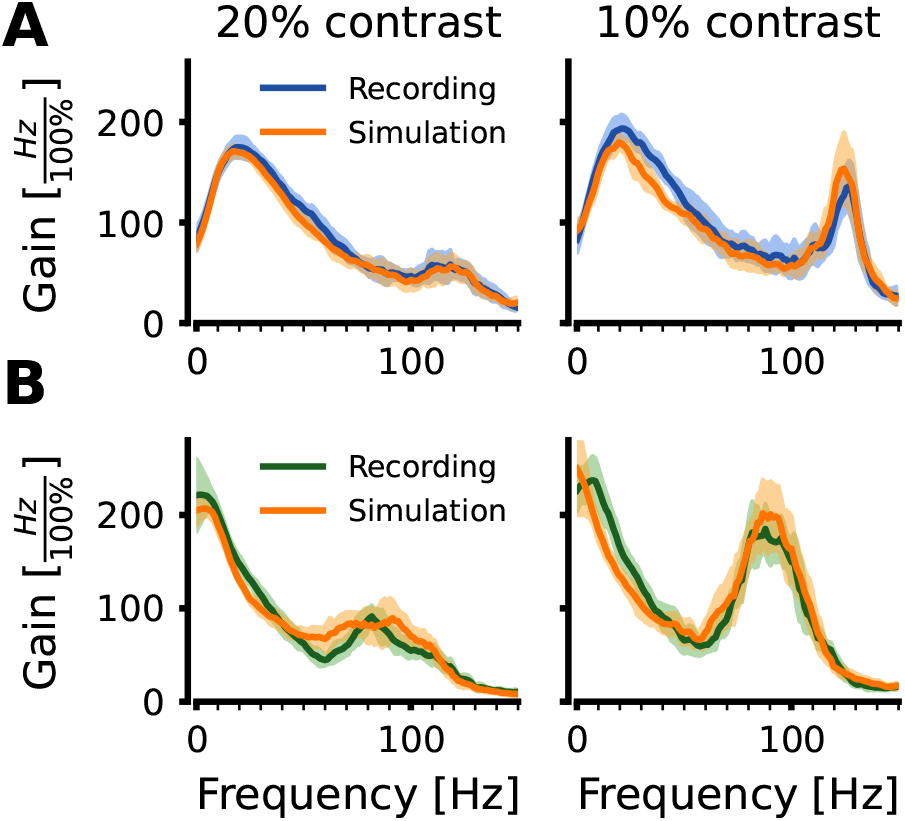
Example gain spectra at the two different contrasts. Comparison of example spectra of *A. leptorhynchus* (**A**) and *E. virescens* (**B**) recordings and simulations at different stimulus intensities (contrast).

**S6 Fig.**
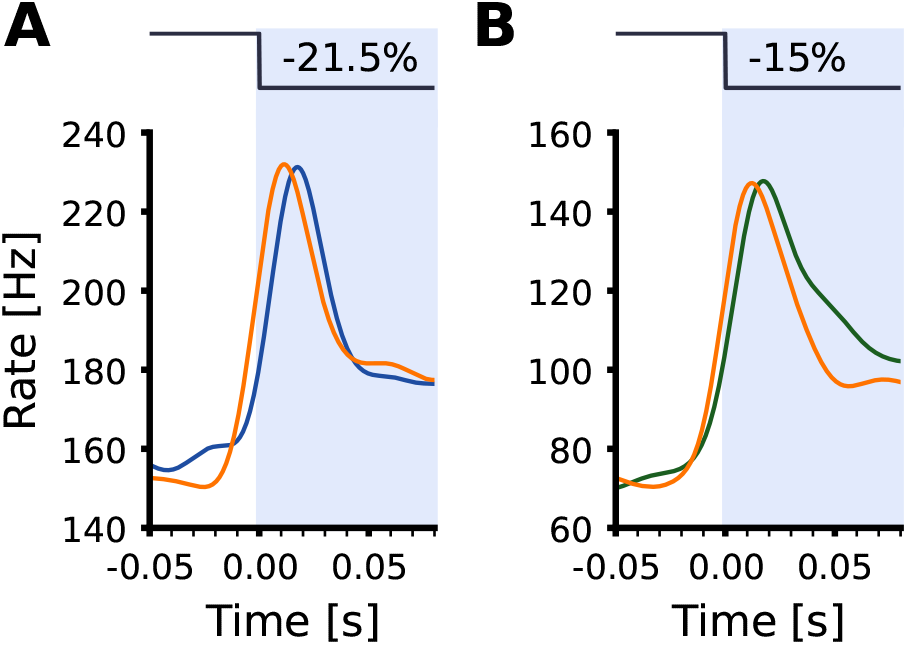
Responses to step stimuli. Comparison of recorded (blue) and simulated (orange) responses to step stimuli. Some experimentally recorded cells were further characterized by their responses to step stimuli. **A** Example of a single step response in *A. leptorhynchus*. Stimulus was a negative step with a contrast of -21.5%. Stimulus onset is at time zero. The size and the time-course of the onset response are well-matched. **B** Same but for an example cell recorded in *E. virescens*. Note that the firing rates were calculated by kernel convolution with an acausal Gaussian kernel of 10 ms that smears out the response that is seems to start before stimulus onset.

## S1 Appendix. Single Ornstein-Uhlenbeck noise is not sufficient

Our model contains two noise sources necessary to reproduce the low-frequency shape of the coherence spectrum (see Fig 9). The white noise added within the adaptation process, *ξ*_*adapt*_, is turned into colored (pink) noise by the adaptation time constant [42]. Here we investigate whether a single Ornstein-Uhlenbeck noise process (*OU-noise* model) which would yield a colored noise directly, would be sufficient to reconstruct the experimental data. For comparison between the original, two-noises, and OU-noise model, we trained two separate SBI networks for the two different models that were based on the same number of training runs to exclude biases based on a larger training set. Both networks were then queried for most-likely models corresponding to each of the recorded cells. We here focus on three features, the serial correlation at lag 1 (*B3: SC*_*ISI*_), the zero-frequency coherence (*C1: C*_0*Hz*_), and the frequency of the maximum coherence (*C4: F*_*max*_).

The 1st-lag serial correlation of the baseline ISI intervals is slightly elevated in the OU-noise models compared to the original models, all models are above the equality line (Fig 10 A, *B3*, gray and orange dots, respectively). The same is true for the zero-Hertz coherence: while the models of the original model are distributed around the equality line, the OU-noise models are consistently above (Fig 10 A, feature *C1*). The frequency of the maximum coherence is most strongly affected and most OU-noise models underestimate the position of the best coherence (Fig 10, feature *C4*). Comparing the goodness of fit of the coherence to the experimental data across all recorded cells, the best models drawn from the SBI-network for the original model perform slightly but not statistically significantly better than the OU-noise models (Fig 10 B). In the following panels of Fig 10 we explicitly selected model cells (from the original model) that show the low-frequency fall-off and tried to resemble these by sampling from the OU-noise network. The OU-noise models all fail to reproduce the initial parts of the coherence spectrum. Even if the peak and the fall-off to higher frequencies are well met (compare red, and dashed red lines in Fig 10 D), the rising part is off. We then perform targeted manipulations of the Ornstein-Uhlenbeck noise to understand the effects of these on the coherence spectrum. In Fig 10 E we change the time-constant of the noise process to change the relative strengths of the low- and high-frequency noise content and compare the resulting spectra to the spectrum of the corresponding original model. Increasing *τ* attenuates the high-frequency content of the noise. As a result, the low-frequency coherence is pushed down but at the same time, the higher frequency coherence is increased due to the improved SNR at higher frequencies (arrows). Using the version that best matches the low-frequency part, and increasing the overall noise strength does not help as the overall stimulus-response coherence is decreased Fig 10 F. We thus conclude that the white noise source in the adaptation process together with the additive current noise in the integrater are needed to resemble the characteristics of the ampullary cells.

**Fig 10.**
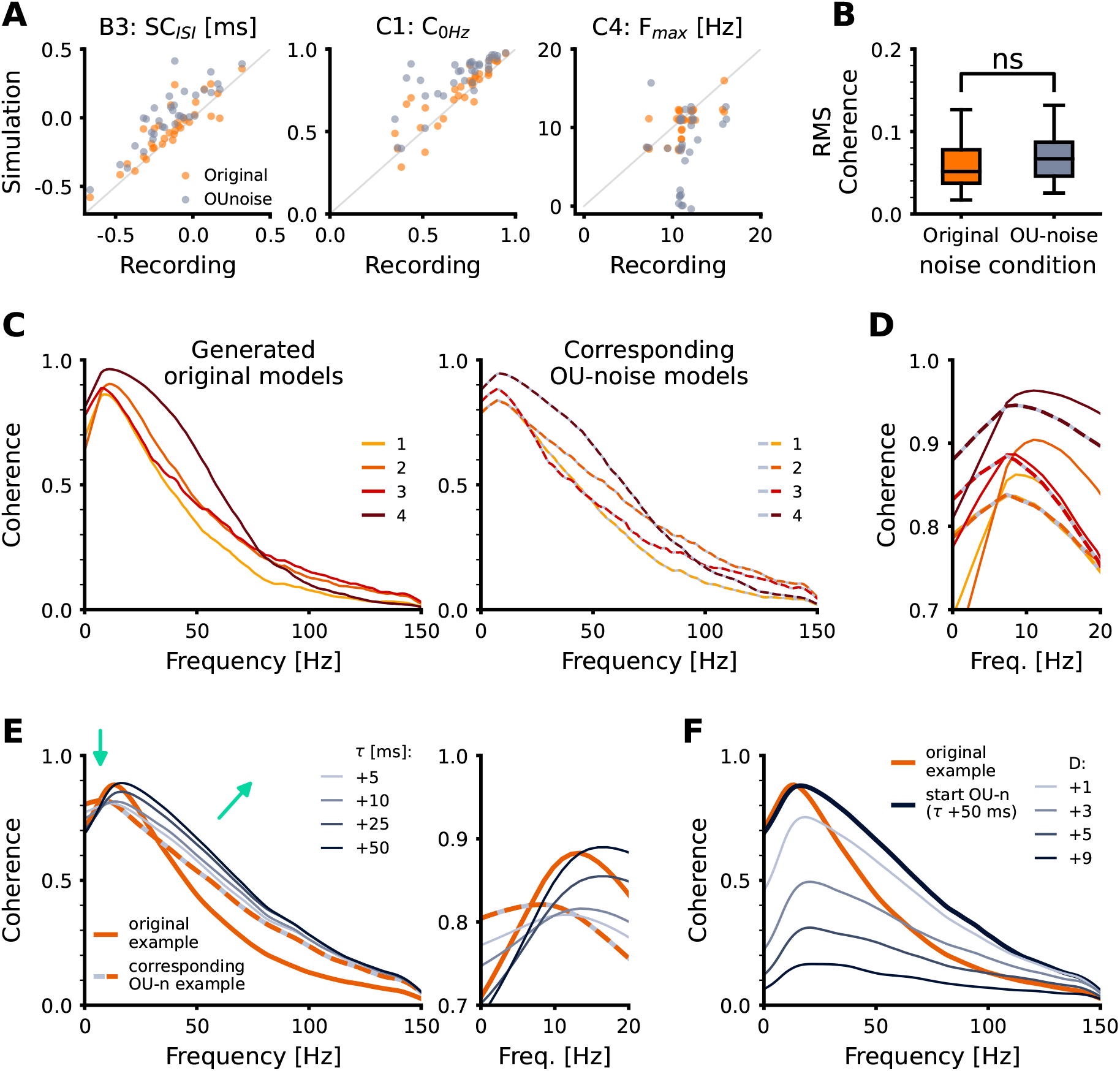
Two noise sources or a single source of colored noise? **A** Comparison of three response features (*B3: SC*_*ISI*_, *C1: C*_0*Hz*_, and *C4: F*_*max*_) of the best models from the original (orange) and the OU-noise model (gray) with the recorded features. **B** Root-mean-square distance between the recorded coherence spectra and the respective spectra of the original and OU-noise models. **C** Coherence spectra of four different original models (left). The respective best matches from the OU-model (right, dashed). **D** Zoom-in on the low-frequency parts of the spectra shown in C. **E** Effect of increased time-constant of the Ornstein-Uhlenbeck process. **F** Attempt to yield a better fit by increasing the overall OU-noise strength fails. Note that we used 100 s of white noise stimulation to achieve smoother spectra.

## S2 Appendix. Application of the trained SBI-network

The trained SBI-network can be applied to draw model parameters that are likely to fit the desired response features best. To make the SBI-network easily accessible, we created a python-written graphical user interface. The Qt-based tool is freely available under the LGPL https://github.com/bendalab/ampullarymodel.

The UI has three main functions, from left to right:

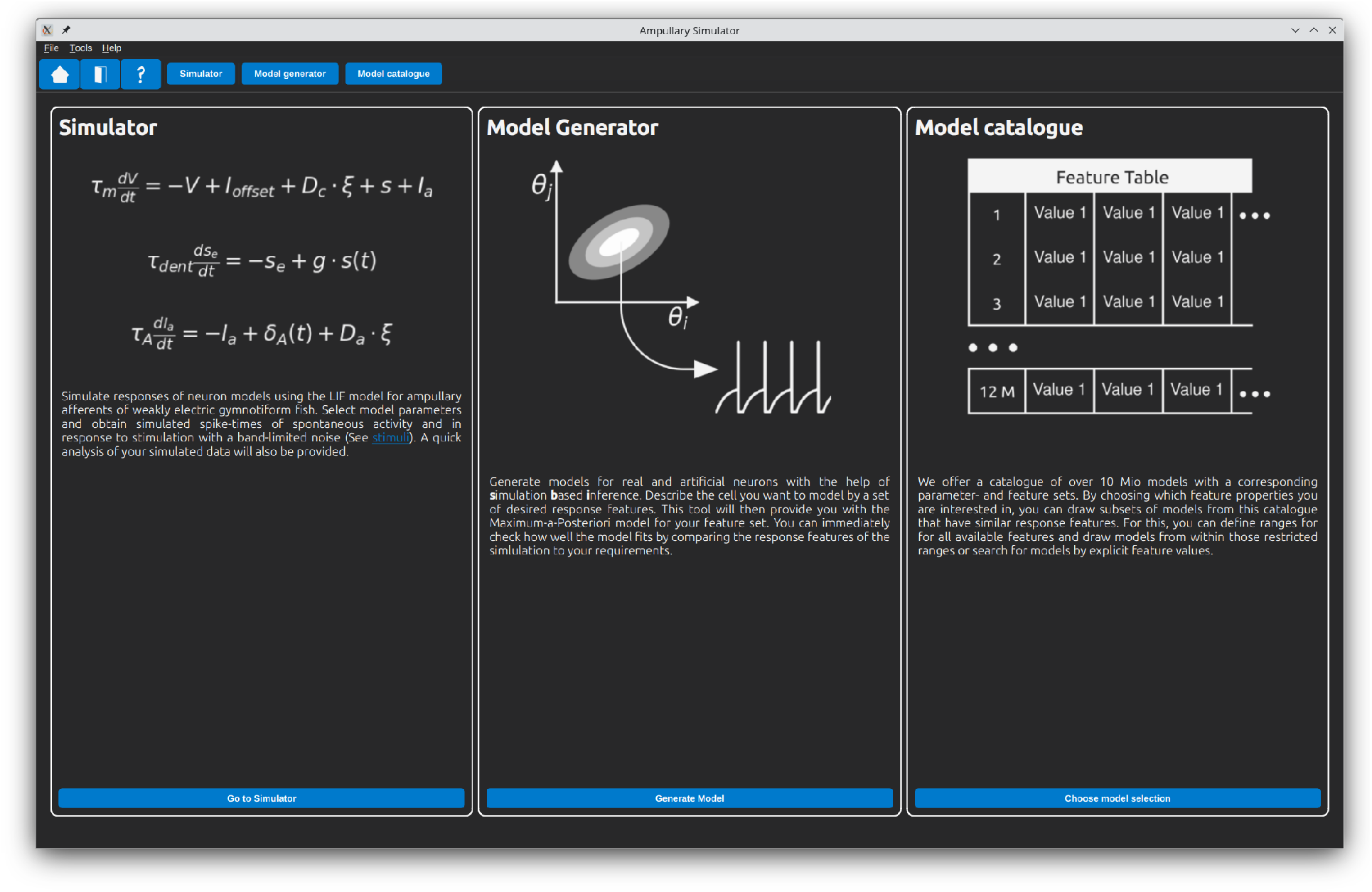

1. Simulator
  - Single cell — Specify model parameters and run a basic simulation. The UI creates a feature overview figure such as Fig 2.
  - Population — Alternatively, provide a csv file with sets of model parameters to run the simulation on a whole population.
2. Model Generator
  - Single cell — Specify response features and retrieve model parameters from the SBI-network for a single cell.
  - Population — Provide a csv file with desired features and retrieve the most-likely model parameters from the SBI-network.
3. Model catalogue
  - Select by range — Specify feature ranges within the bounds of the training set and retrieve sets of random models that fall into the selected feature ranges.
  - Select by feature — Explicitly specify selected features and retrieve sets of models that try to match the specification but are free in other features.

Of course the network can also be programmatically employed to retrieve model parameters. For this, one needs to install the SBI package https://sbi.readthedocs.io/en/stable/ together with its dependencies and to download the trained network which is available under https://gin.g-node.org/jgrewe/ampullary_sbi/src/master/packages. The following minimal-example shows how to use it:

~~~
import pickle
from pathlib import Path
# load the posterior from file --> type(posterior) = sbi.Posterior
path = Path.cwd() / “apteronotus_ampullary_posterior_1.0.pkl”
with open(path, ‘rb’) as handle:
 posterior = pickle.load(handle)
# initialize the posterior with the desired features and get the mapped posterior
features = np.array([…])
p = posterior.set_default_x(features)
mapped_posterior = p.map(num_iter=1000, show_progress_bars=False)
model_params = mapped_posterior.tolist()[0]
~~~

For historical reasons, the features are ordered as follows. The sequence deviates slightly from the more natural ordering used in this paper:

~~~
[B1, B2, B3, R1, C5, C3, C4, C2, C1, G1, G2, G4, G6, G7, G3, G5, G8], see Fig 2 for reference.
~~~

